# Unconventional conservation reveals structure-function relationships in the synaptonemal complex

**DOI:** 10.1101/2021.06.16.448737

**Authors:** Lisa E. Kursel, Henry D. Cope, Ofer Rog

## Abstract

Functional requirements constrain protein evolution, commonly manifesting in conserved primary amino acid sequence. Here, we extend this idea to secondary structural features by tracking their conservation in essential meiotic proteins with highly diverged sequences. The synaptonemal complex (SC) aligns parental chromosome pairs and regulates exchanges between them. In electron micrographs of meiocytes from all eukaryotic clades, the SC appears as a ~100 nm-wide ladder-like structure with regular striations. Despite the conserved ultrastructure and functions, the proteins that make up the SC are highly divergent in sequence. Here we found that, within the *Caenorhabditis* genus, SC proteins are significantly more diverged than other proteins. However, SC proteins have highly conserved protein length and coiled-coil domain structure. The same unconventional conservation signature holds true for SC proteins in *Drosophila* and mammals, suggesting it could be a universal feature of SC proteins. We used this evolutionary signature to identify a novel SC protein in the nematode *Pristionchus pacificus*, Ppa-SYP-1, which has no significant homology to any protein outside of *Pristionchus*. Our work suggests that the length and relative arrangement of coiled-coils play a key role in the structure and function of the SC. Furthermore, our analysis implies that expanding sequence analysis beyond measures of per-site identity or similarity can enhance our understanding of protein evolution and function.

**Short abstract:** Functional requirements constrain protein evolution, commonly manifesting in a conserved amino acid sequence. Here, we extend this idea to secondary structural features by tracking their conservation in essential meiotic proteins with highly diverged sequences. The synaptonemal complex (SC) is a ~100 nm-wide ladder-like meiotic structure present in all eukaryotic clades, where it aligns parental chromosomes and regulates exchanges between them. Despite the conserved ultrastructure and functions of the SC, SC proteins are highly divergent within *Caenorhabditis*. However, SC proteins have highly conserved length and coiled-coil domain structure. We found the same unconventional conservation signature in *Drosophila* and mammals, and used it to identify a novel SC protein in *Pristionchus pacificus*, Ppa-SYP-1. Our work suggests that coiled-coils play wide-ranging roles in the structure and function of the SC, and more broadly, that expanding sequence analysis beyond measures of per-site similarity can enhance our understanding of protein evolution and function.

## Introduction

Functional and structural constraints leave evolutionary signatures on proteins. Often, functionally important domains undergo purifying selection and tend to evolve slowly. For example, enzymatic active sites require precise positioning of amino acids and can be identified based on sequence conservation. Even seeming exceptions are telling. Many genes in the immune system are fast-evolving and undergo recurrent changes (positive selection) to uphold tight interaction interfaces with foreign proteins (1–3).

This paradigm of protein evolution holds true for most studied proteins, but several exceptions have been identified (4, 5). One such example is the protein components of the synaptonemal complex (SC), and specifically, the central region of the SC (referred to throughout as ‘the SC’; Fig. 1A). The SC is present in all eukaryotic clades, and is essential for meiosis. It brings parental chromosomes into close proximity in meiotic prophase and forms the interface between them. The SC also regulates genetic exchanges (crossovers), which serve as the physical link between parental chromosomes during the first meiotic division. First observed more than 60 years ago (6, 7), the SC is a 100 nm wide, ladder-like structure with regularly-spaced rungs (Fig. 1A). In the decades since, electron microscopy allowed the characterization of the SC in meiocytes from numerous organisms, where its ultrastructure was found to be remarkably conserved (8–12). The advent of molecular genetics allowed cloning of SC proteins and revealed that they are perplexingly divergent and cannot be identified across distant taxa based on sequence homology. Despite poor per-site identity, in cases where SC proteins have been cloned at least one assumes a stereotypical head-to-head orientation spanning the space between the parental chromosomes: N-termini pointing toward the center of the SC and C-termini pointing outwards (Fig. 1A). SC proteins with this orientation are referred to as ‘transverse filaments’ and help determine the width of the SC via a central coiled-coil domain (13–16). Thus, SC proteins represent a case where functional and ultrastructural conservation does not seem to constrain a protein’s primary sequence – a pattern that, although unconventional for essential proteins, is likely to be more prevalent than is currently appreciated (17).

**Fig. 1:**
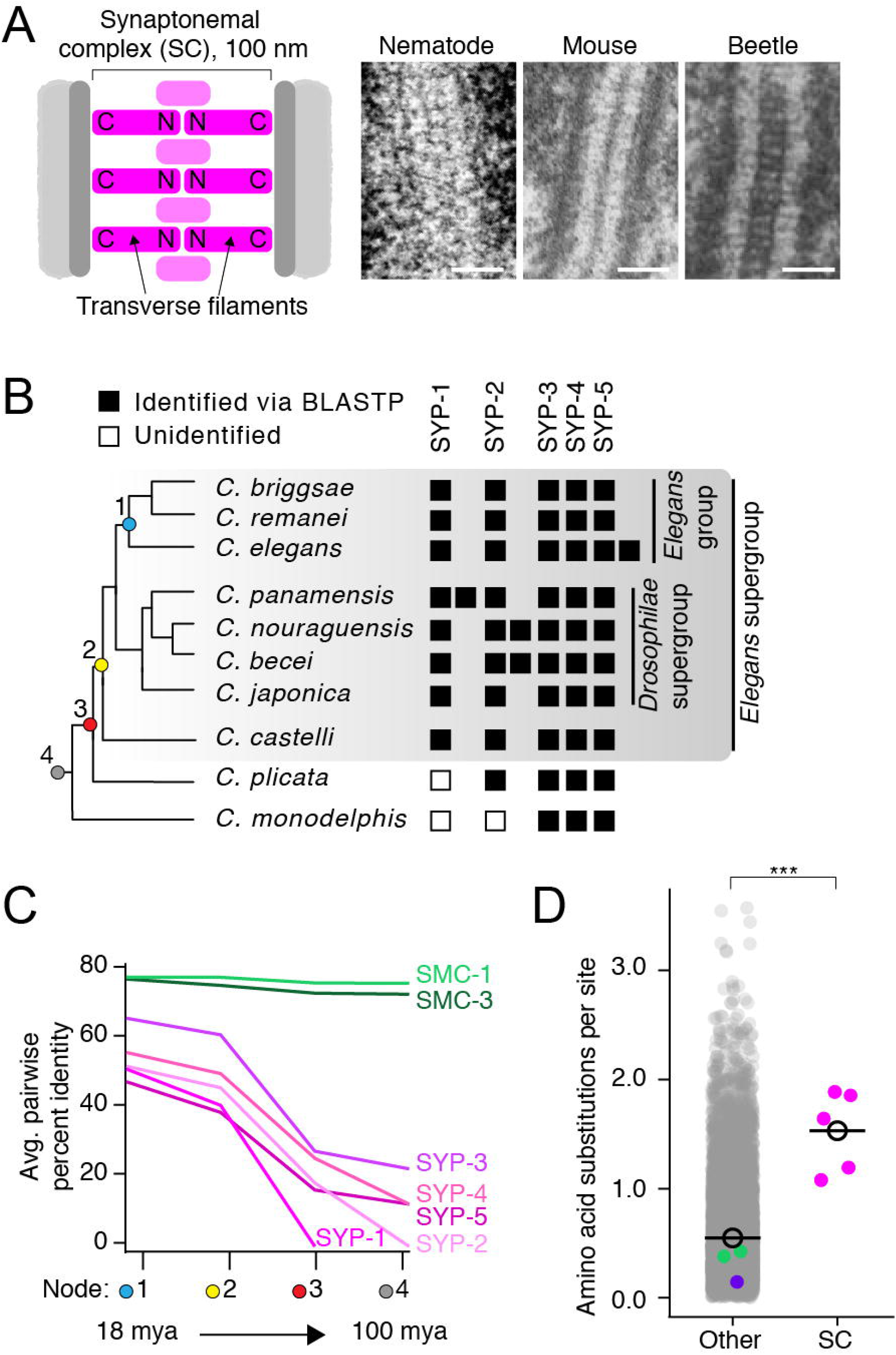
The divergence of SC proteins is driven by neutral evolution. (A) Left, diagram of the SC. The N- and C-termini of transverse filaments are labeled. SC = pink, axial element (scaffold for SC assembly) = dark grey, chromosomes = light grey. Right, electron micrographs of the SC from nematode (*C. elegans*), mouse (*M. musculus*) and beetles (*B. cribrosa*), demonstrating the conserved organization and dimensions of the SC (adapted from (11, 46, 76)). Scale bars = 100 nm. (B). Abridged *Caenorhabditis* species tree. Presence of SC proteins is to the right of each species. Filled box = present, unfilled box = no ortholog identified. (C) Graph of average pairwise percent identity for SC proteins and SMC-1/3 (two chromatin associated coiled-coil proteins) as controls. Colored nodes on the x-axis correspond to the species tree in (B). Evolutionary time increases from left to right with time estimates according to (24) listed below select nodes. (D) Dot plot comparing amino acid substitutions per site of SC proteins to all other *Caenorhabditis* proteins. Black circle = median value. The median SC amino acid substitutions per site = 1.64, other = 0.43, p-value = 0.0005. SMC-1/3 (cyan) and LEV-11 (purple) are highlighted as controls. The high divergence of SC proteins cannot be explained by positive selection (Table S1).

Here we attempt to resolve this paradox of SC protein evolution. We find that SC proteins in *Caenorhabditis*, although highly divergent, harbor features that are strikingly conserved: protein length, as well as the length and location of the coiled-coils. The conservation of these features is also exhibited by SC proteins in *Drosophila* and *Eutherian* mammals. We harness this intra-clade evolutionary signature to predict and identify a novel SC protein in the nematode *Pristionchus pacificus*. Our findings suggest that the fitness landscape of SC proteins is governed by secondary structures, shedding light on structure/function relationships of this conserved chromosomal interface.

## Results

### SC proteins are ancient and preserved in Caenorhabditis

To analyze the evolution of SC proteins, we generated and refined a dataset of all known SC proteins from 25 *Caenorhabditis* species. These species, many of which have been sequenced in the last two years, represent the *Elegans* and *Drosophilae* supergroups as well as two basally branching *Caenorhabditids*, *C. plicata* and *C. monodelphis* (Fig. 1B, Fig. S1). In the model organism *C. elegans,* six SC proteins have been identified: SYP-1 – SYP-6 (10, 18–21). SYP-1, SYP-5 and the *C. elegans-*specific SYP-5 paralog, SYP-6, are transverse filament proteins (10, 18, 19, 22, 23). All SC proteins are present in the *Elegans* and *Drosophilae* supergroups, indicating they have been preserved for over 30 million years (24) (Fig. 1B, Fig. S1). This broad preservation is not surprising given that SC proteins in *C. elegans* are inter-dependent for their function and that their elimination causes a dramatic drop in viable progeny (10, 18–21). In addition, we found three instances of protein duplication; the previously identified paralogs SYP-5 and SYP-6 in *C. elegans* (18), a SYP-1 duplication in *C. panamensis* and a SYP-2 duplication in the common ancestor of *C. nouraguensis* and *C. becei* (Fig. 1B, Fig. S1, S2).

To identify SC proteins in the early diverging *Caenorhabditis* species, *C. plicata* and *C. monodelphis,* we used all of our previously identified sequences as queries in BLASTP and tBLASTn searches, with a lenient e-value cutoff (1.0e^−1^). This allowed us to identify SYP-3, −4 and −5 orthologs in both *C. plicata* and *C. monodelphis* (Fig. 1B, Fig. S1, S2). We also found SYP-2 in *C. plicata* but not in *C. monodelphis.* We were unable to identify SYP-1 in either *C. plicata* or *C. monodelphis* (Fig. 1B, Fig. S1). It is possible that *C. plicata* and *C. monodelphis* have fewer SC proteins. However, given the fact that SC proteins are essential for meiosis and functionally inter-dependent, a plausible hypothesis is that the SC proteins are too diverged to be detected in these distantly related species. In line with this possibility, SC protein amino acid percent identity drops rapidly as more distantly related species are included in the comparison (Fig. 1C).

### Neutral evolution drives the high divergence of Caenorhabditis SC proteins

The difficulty in identifying SC proteins in distantly related *Caenorhabditis* species is not surprising. *Drosophila* SC proteins were not found in other insects (25). Among vertebrates, sequence similarity of the transverse filament protein SYCP1 is limited to two short motifs, which are absent in *Caenorhabditis* and *Drosophila* (26). We found that SC proteins are among the most diverged proteins in *Caenorhabditis*. On average, there are significantly more amino acid substitutions per site in SYP-1 – SYP-5 compared to the *Caenorhabditis* proteome (SC proteins: median amino acid substitutions per site = 1.64, other proteins = 0.43, p-value = 0.0005, Fig. 1D). This divergence is not merely due to the prevalence of coiled-coils: the sequences of SMC-1, SMC-3, and LEV-11 which harbor extensive coiled-coils, are highly conserved in *Caenorhabditis* (Fig. 1C-D, see also (27)).

The role of the SC in regulating genetic exchanges, and consequently chromosome segregation, makes it a candidate for involvement in meiotic drive, where a genetic locus skews its own inheritance. Meiotic drive often incurs a fitness cost, creating pressure for the emergence of suppressors. This tit-for-tat evolutionary arms race leads to rapid evolution, which can be detected bioinformatically as positive selection. Indeed, meiotic drive has been invoked to explain the rapid evolution of SC proteins in *Drosophila* (25). However, we found no evidence for positive selection in any *Caenorhabditis* SC protein (Table S1). Using the CodeML program from PAML (28), we found no significant difference between models M8a (no positive selection allowed, dN/dS <= 1) and model M8 (positive selection allowed). Consistent with the high divergence observed above (Fig. 1C-D), we found that fewer than 50% of sites evolve under purifying selection when examined on a per-site basis using a Fixed Effects Likelihood model (Table S1). Our per-site analysis found almost no evidence of positive selection (no sites in SYP-1 or SYP-2 and only two sites each in SYP-3, −4 and −5; Table S1, Fig. S3A). Altogether, our analysis indicates that neutral evolution (lack of constraint) explains the high divergence of SC proteins in the *Caenorhabditis* lineage.

### Protein length and coiled-coil domains are conserved in SC proteins

Despite the poor conservation of primary amino acid sequence in SC proteins (Fig. 1, Table S1 and (25)), bioinformatic and functional analysis has pointed to the prevalence of coiled-coils (29). We used Paircoil2 to predict the likelihood that each position is part of a coiled-coil (plotted throughout as a “coiled-coil score” [1 – Paircoil2 score]; (30); Fig. 2A). These plots reveal striking conservation of the position and length of coiled-coils.

**Fig. 2:**
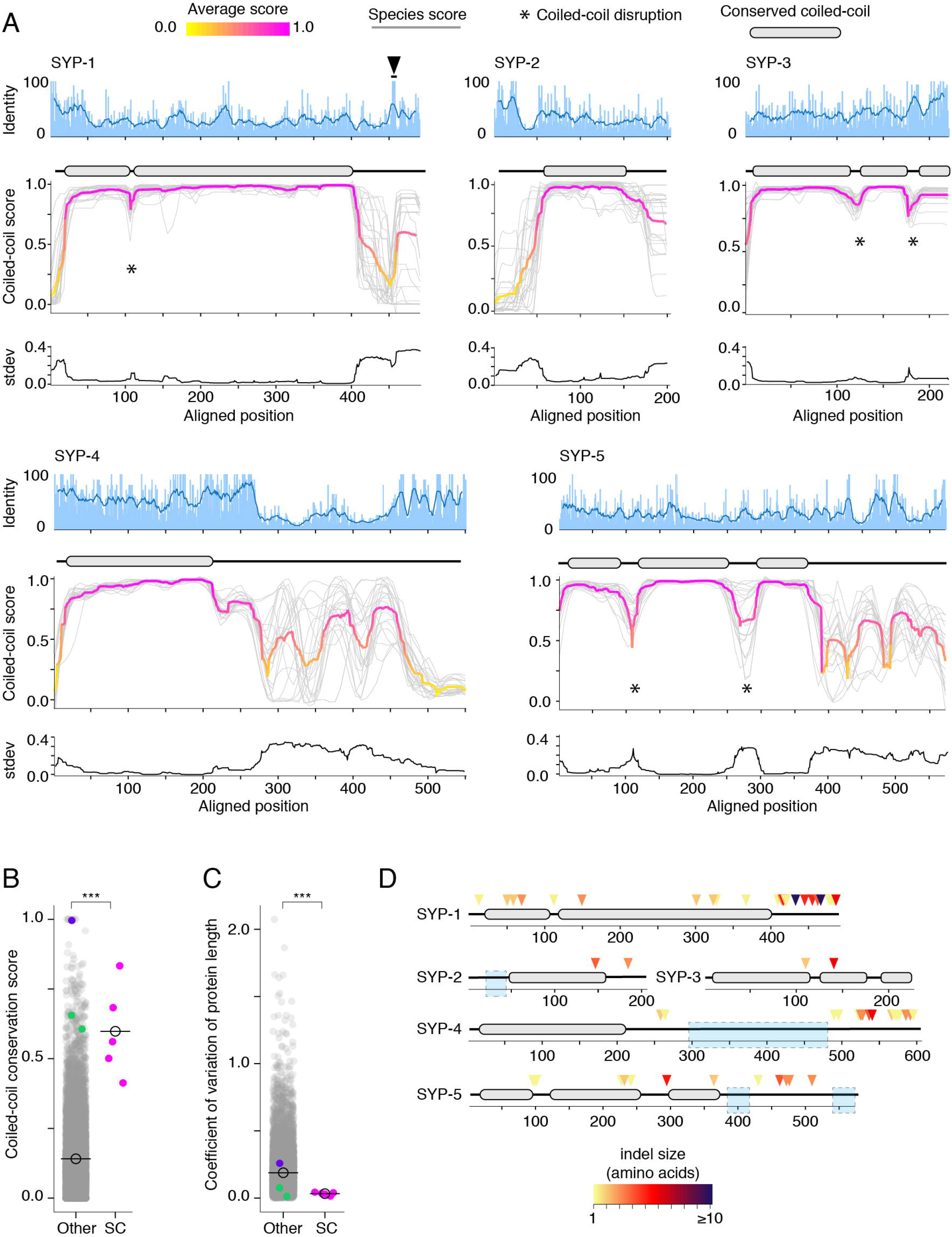
Protein length and coiled-coil domains are conserved features of SC proteins. (A) Percent amino acid identity (top), coiled-coil score (middle) and standard deviation of coiled-coil score (bottom) at each aligned position for SYP-1 – SYP-5. Sliding window average of percent identity is shown (blue line). Coiled-coil conservation plots display the coiled-coil score (1 – Paircoil2 score) at each aligned position for all SYP proteins from each species (grey lines). Magenta and yellow line = average score at each position. Gene models depicting conserved coiled-coil domains (grey-filled ovals) are shown above each coiled-coil conservation plot. The black arrowhead indicates the conserved Polo-box Domain in SYP-1. Note that coiled-coil conservation is generally not reflected in elevated (or diminished) amino acid identity with the exception of SYP-4. (B) Dot plot comparing the coiled-coil conservation score of SC proteins to all other proteins in *Caenorhabditis.* The coiled-coil conservation score is the average minimum value of the coiled-coil score (1 – Paircoil2 score) for each position. Median coiled-coil conservation score for SC proteins = 0.56, median for all other proteins = 0.12. p-value = 0.00019. (C) Dot plot comparing coefficient of variation of protein length of SC proteins to all other proteins in *Caenorhabditis*. Median for SC proteins = 0.04, median for all other proteins = 0.16. p-value = 0.0004. SMC-1/3 (cyan) and LEV-11 (purple) are shown as controls in (B) and (C). LEV-11 (tropomyosin) coiled-coil conservation score of 0.996 is consistent with the importance of coiled-coils for its function (77). (D) Indels in Elegans supergroup SC proteins. Gene model of SYP-1 – SYP-5 coiled-coil domains with indel positions marked with colored arrowheads, with darker reds indicating larger indels. Light blue boxes surround regions that were excluded from analysis due to alignment uncertainty.

For example, in SYP-1, the coiled-coil begins precisely at position 40 and ends at position 400 in all species (Fig. 2A). Disruptions, observable as dips in an otherwise continuous coiled-coil, are also conserved (marked by asterisks in Fig. 2A). SYP-1 has a short (<10 amino acid) disruption at position 110 whereas SYP-3 and SYP-5 have longer disruptions (20 – 50 amino acids). These disruptions might create bends in the otherwise rod-like structure of SC proteins (31), similar to the conserved “kinks” and “elbows” in the coiled-coils of the kinetochore protein NDC80 (32, 33) and the ring-like SMC-family proteins (34, 35).

To quantitate the extent of coiled-coil conservation, we developed a scalar metric, the coiled-coil conservation score, that takes the minimum score (least likely to be part of a coiled-coil) from every aligned position (as in Fig. 2A). This score is averaged across the alignment. Proteins with coiled-coils in the same position will have a higher score than proteins whose coiled-coils do not overlap or that lack extended coiled-coils altogether. Consistent with our qualitative observations, SC proteins have a significantly higher coiled-coil conservation scores on average compared to other proteins (Fig. 2B; median coiled-coil conservation score for SC proteins = 0.56, other = 0.12, p-value = 0.00019). This stands in contrast to their higher-than-average amino acid divergence (Fig. 1D). Neither coiled-coils nor their edges leave discernable signatures of sequence conservation on SC proteins (Fig. 2A). We also do not find strong correlation between the coiled-coils and amino acids under purifying selection (Fig. S3A). These observations likely reflect the relatively loose requirements for coiled-coil formation: heptad repeats where the first and fourth amino acids are hydrophobic and the fifth and seventh amino acids are charged or polar (Fig. S3B; (36)).

We wondered whether other secondary structural features are conserved in SC proteins. SC proteins often encode disordered domains of unknown function (37). We used PONDR VL3 to predict the likelihood of disorder for all sites in *Caenorhabditis* SC proteins (Fig. S4). Unlike coiled-coils, the length and position of disordered domains was mostly varied between species. However, a few disordered regions were conserved, including the C-termini of SYP-1, −3, −4 and −5 and the N-terminus of SYP-2 (Fig. S4). This analysis indicates that while multiple secondary structures might be under selection in SC proteins, conservation is particularly strong for coiled-coils.

Finally, we explored whether the conservation of coiled-coils in SC proteins is reflected in their overall length. We analyzed coefficient of variation of protein length of all *Caenorhabditis* proteins. We found that the median variation in length of SC proteins is significantly lower than that of other *Caenorhabditis* proteins (Fig. 2C, median coefficient of variation of protein length for SC proteins = 0.03, other = 0.16, p-value = 0.0004), again, in striking contrast to their diverged primary amino acid sequence (Fig. 1C, 1D). Low variation of protein length suggests strong purifying selection against insertions and deletions (indels). Indeed, we find a significant depletion of indels in the coiled-coils of SYP-1 through SYP-5 compared with regions outside the coiled-coils (Fig. 2D, Table S2; two-tailed p-value from Fisher’s test comparing alignment positions with indels to alignment positions that are part of coiled-coils < 0.0001), consistent with the conservation of their length and arrangement.

Taken together, these analyses highlight several conserved features that are not apparent from primary amino acid conservation alone. We find that SC proteins show an unusual evolutionary signature consisting of three key features: (1) high amino acid divergence, (2) conserved coiled-coils and (3) low coefficient of variation of protein length. To demonstrate this point, we plotted these metrics for all proteins in *Caenorhabditis* on a Cartesian coordinate system (Fig. 3A, S5A). While most proteins form a pyramidal cluster near the origin, SC proteins are among the few proteins situated away from it (Fig. 3A, S5A). Of the few proteins clustering with SC proteins are several that play a role in the mitotic spindle (Fig. S5B, S5C; p-value = 0.0076). Noteworthy among them is SPD-5, a component of the pericentrosomal material that nucleates spindle microtubules (Fig. S5D; (38)). Like SC proteins, SPD-5 and its apparent functional homologs in other eukaryotic clades - Cdk5Rap2 in vertebrates and Centrosomin in Drosophila – do not share significant sequence homology despite their conserved and essential functions in cell division.

**Fig. 3:**
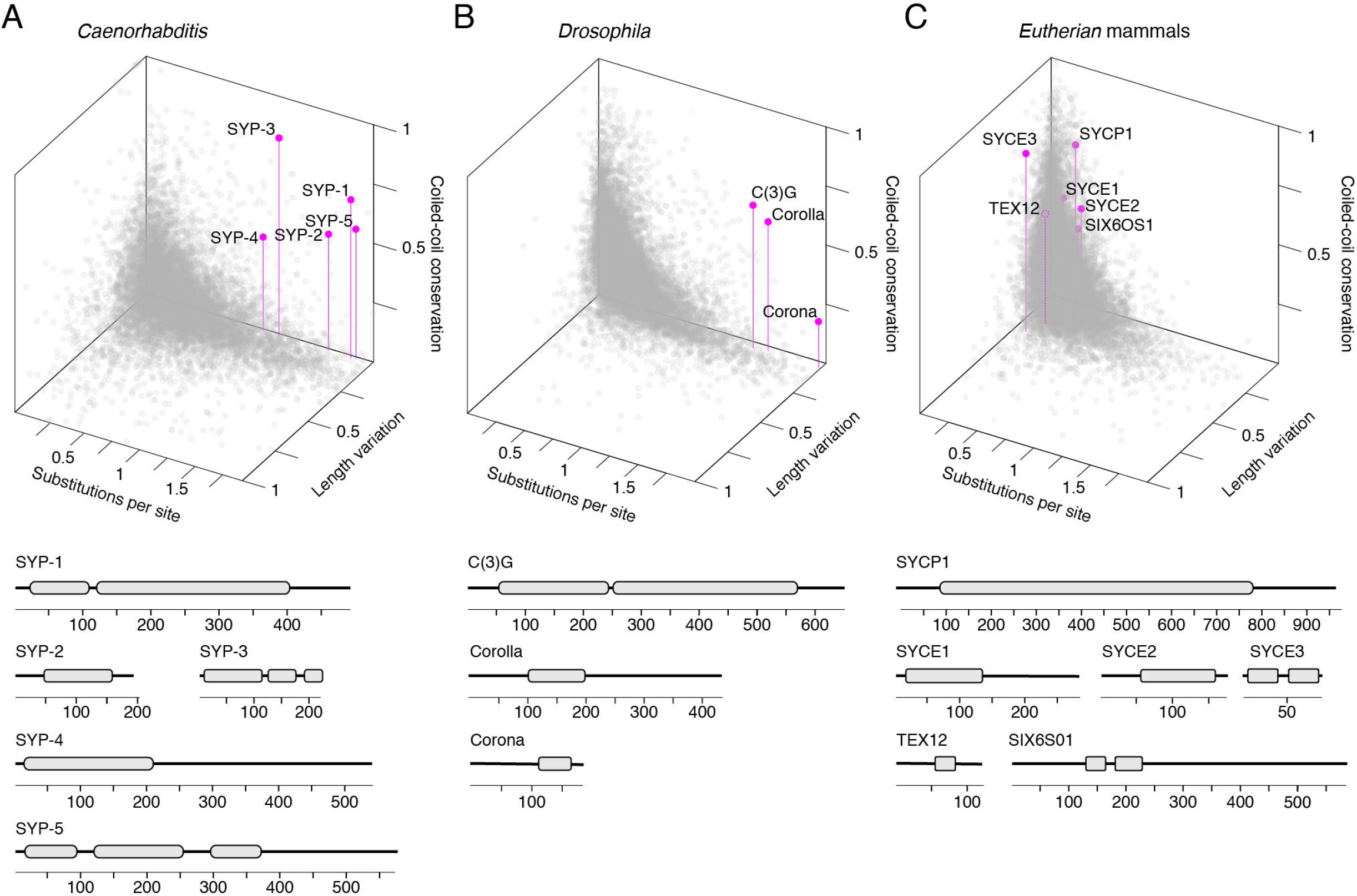
SC proteins have an unconventional, but conserved, evolutionary signature. 3D scatter plot comparing amino acid substitutions per site, coefficient of variation of protein length and coiled-coil conservation score of all proteins in 25 *Caenorhabditis* species (A), 30 *Drosophila* species (B) and 15 mammalian species (C). SC proteins = pink dots with vertical lines, other proteins = grey dots. Gene models of SC proteins depicting conserved coiled-coils derived from conservation plots (Fig. 2, S6 and S7) are shown below each scatter plot.

### The evolutionary signature of SC proteins is conserved across phyla

Next, we wondered whether this evolutionary signature was restricted to SC proteins in *Caenorhabditis*. In a dataset of 30 *Drosophila* species spanning 40 million years of evolution, the three known *Drosophila* SC proteins – Corolla, Corona and the transverse filament protein C(3)G - exhibit a similar evolutionary signature to SC proteins in *Caenorhabditis*, and likewise, occupy a region of the coordinate system occupied by few other proteins (Fig. 3B, S6).

Analysis of proteomes of 15 mammalian species representing Xenarthra, Afrotheria, Laurasiatheria and Euarchontoglires (*Eutherian* mammals, ~100 million years) revealed a similar, albeit weaker, evolutionary signature of the six SC proteins SYCP1, SYCE1-3, TEX12 and SIX6OS1 (Fig. 3C, S7). While mammalian SC proteins exhibited conserved coiled-coil domains and low variation in protein length, they had a lower overall divergence compared to *Caenorhabditis* and *Drosophila* SC proteins (median amino acid substitutions per site for mammalian SC proteins = 0.26, compared to 1.64 in *Caenorhabditis* and 1.69 in *Drosophila)*. This might be explained by the overall lower median divergence of the proteome along the mammalian lineage. When their divergence is ranked against the rest of the proteome, *Caenorhabditis* and *Drosophila* SC proteins are all in the 96^th^ percentile or above. Mammalian SC protein divergence ranges from 42^nd^ (SYCE3) to 91^st^ percentile (SYCE2), with four out of six proteins (SYCP1, SYCE1, SYCE2 and SIX60S1) ranking above the 67^th^ percentile. The relatively constrained divergence of mammalian SC proteins, including along their coiled-coils, might indicate novel functions adopted by these domains in mammals. In addition, N- and C-terminal extensions in *Loxodonta africana* (African elephant) SYCE3, caused this protein to exhibit an overall higher coefficient of variation of length. Despite these differences, the shared evolutionary signatures of SC proteins in mammals, *Drosophila* and *Caenorhabditis* suggests that these conserved features reflect structural and/or functional constraints acting on the SC.

### Identification of a novel SC protein in Pristionchus pacificus

Most SC proteins have been identified independently in each lineage by genetic and cell-biological methods (10, 18–21, 39–41). The strong evolutionary signature of SC proteins raised the possibility that we could identify SC proteins *in silico* by relying on intra-clade conservation patterns. To test this, we turned to the nematode genus *Pristionchus,* which is distantly related to *Caenorhabditis*. While *Pristionchus pacificus* is an emerging model organism for evolutionary and developmental biology, its SC is poorly characterized. One SC protein, Ppa-SYP-4, has been identified based on a limited sequence similarity to the C-terminus of *C. elegans* SYP-4 (42). Notably, Ppa-SYP-4’s coiled-coil is predicted to be only 31nm long, too short to span the 100nm SC as a head-to-head dimer, suggesting that we have yet to identify a transverse filament protein in *P. pacificus*. We developed a bioinformatics pipeline to identify SC proteins based on our prior analysis of *Caenorhabditis, Drosophila* and mammals. Rather than leveraging sequence homology across distant genera, we categorized the proteome of eight sequenced *Pristionchus* species based on our evolutionary signature – high amino acid substitutions per site, low coefficient of variation of protein length and high coiled-coil conservation scores – and generated a candidate list of *Pristionchus* SC proteins (Fig. 4A, 4B). We further filtered our list for germline enriched genes (see Methods) and were left with only eight candidate SC proteins, one of which was, gratifyingly, Ppa-SYP-4.

**Fig. 4:**
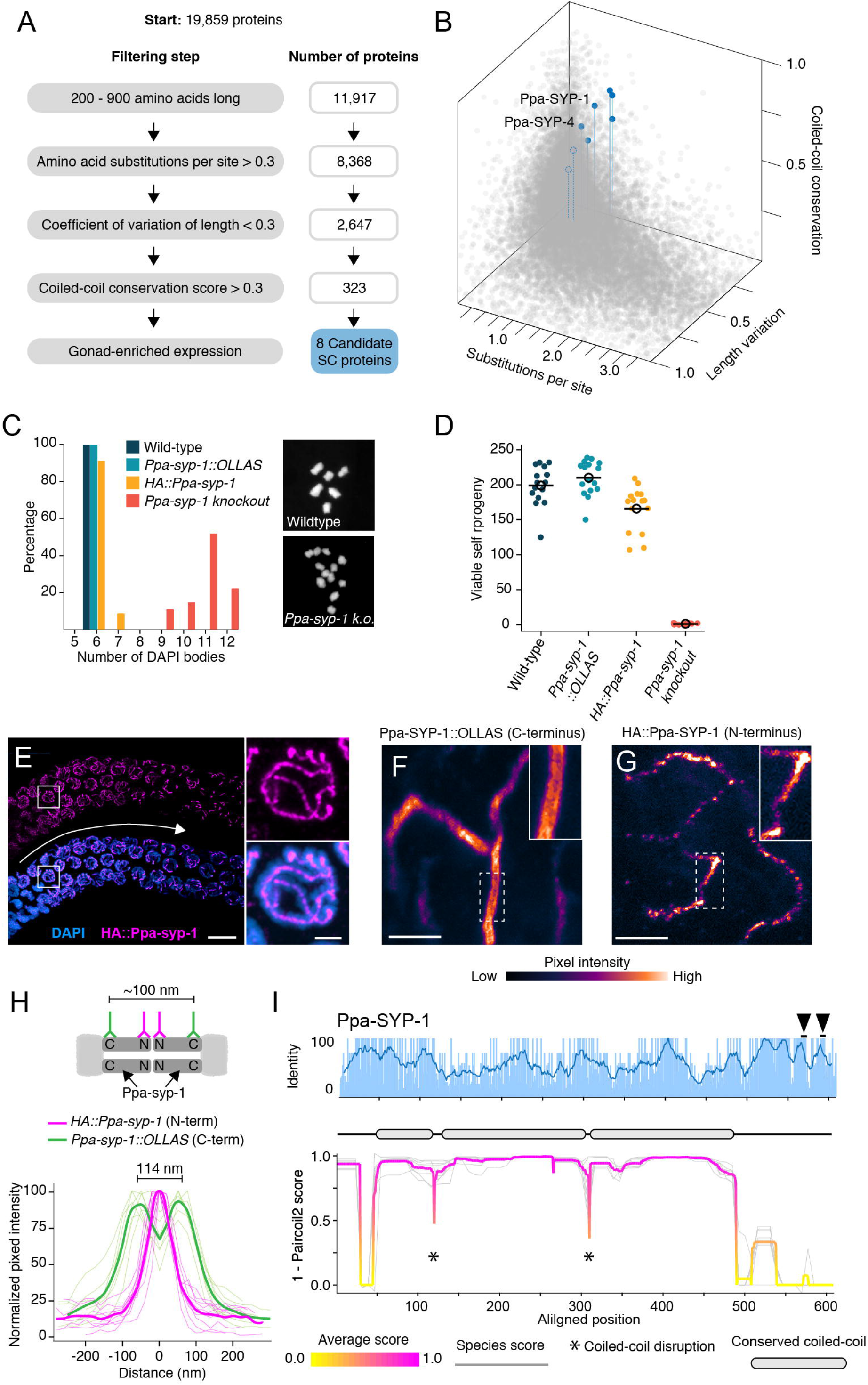
Identification of *P. pacificus* SYP-1. (A) Flow chart of filtering steps to identify candidate SC proteins in *Pristionchus*, with the number of remaining proteins after each step shown to the right. See Methods for details. (B) 3D scatter plot comparing amino acid substitutions per site, coefficient of variation of protein length and coiled-coil conservation score of all *Pristionchus* proteins. Blue dots = candidate SC proteins, other = grey dots. (C-D) Number of DAPI bodies (C) and total viable self-progeny (D) from wild-type *P. pacificus, Ppa-syp-1::OLLAS, HA::Ppa-syp-1,* and *Ppa-syp-1* knockout. In (C), representative DAPI body images are shown for wild-type and *Ppa-syp-1* knockout. (E) Left, image of a prophase region of a gonad from a *HA::Ppa-syp-1* hermaphrodite where meiosis progresses from left to right (white arrow), stained with antibodies against HA (magenta). Scale bar = 5 μm, scale bar in inset = 2 μm. (F - G) STED images of a representative chromosome from *Ppa-syp-1::OLLAS* (F, C-terminus) and *HA::Ppa-syp-1* (G, N-terminus) hermaphrodites stained with antibodies against OLLAS or HA tags, respectively (colored according to pixel intensity). Insets show higher magnification of sections in the dashed boxes. Note the ‘railroad tracks’ configuration in (F). Scale bars in (F – G) = 1 μm. (H) Top, model depicting Ppa-SYP-1 as a transverse filament protein, with antibodies targeting the C- and N-termini. Expected distance between C-terminal epitopes is ~100 nm. N-terminal epitopes are expected to be too close to be resolved. Bottom, line scans of normalized pixel intensity across the SC in *Ppa-syp-1::OLLAS* (green) and *HA::Ppa-syp-1* (magenta). Bold green and magenta lines represent the average of multiple line scans. (I) Average pairwise percent amino acid identity (top) and coiled-coil score (bottom) for Ppa-SYP-1 from eight *Pristionchus* species. Sliding window average of percent identity is shown (blue line). Observable steps in the percent identity bar graphs are attributed to fewer species in the *Pristionchus* proteome dataset (8 *Pristionchus* species vs. 25 *Caenorhabditis* species in Fig. 2). Magenta and yellow line in coiled-coil plot = average score at each position, grey lines = individual species scores. Gene model depicting conserved coiled-coil domains (grey-filled ovals) is shown above the coiled-coil plot. Black arrowheads point to two Polo-box Domains in Ppa-SYP-1, which are conserved in *Pristionchus*.

We generated null alleles in our top three candidates using CRISPR/Cas9, and found that one of these genes, PPA16075, is an SC protein that we named Ppa-SYP-1. Examination of condensed meiotic chromosomes (DAPI bodies) revealed that almost all chromosomes in *Ppa-syp-1* lacked crossovers (Fig. 4C). Consistent with this, *Ppa-syp-1* mutant hermaphrodites give rise to almost no viable self-progeny; a reflection of embryonic aneuploidy caused by uncoordinated chromosome segregation during meiosis (Fig. 4D). To visualize Ppa-SYP-1, we generated two strains with internal tags: an HA tag near the N-terminus and an OLLAS tag near the C-terminus, with only minor effects on SC function (Fig. 4C, 4D, and Methods). As expected for an SC protein, HA::Ppa-SYP-1 localized to the interface between the parental chromosomes in meiotic prophase (Fig. 4E). Its C-terminus formed parallel tracks spaced 114 nm apart on average whereas the N-terminus formed a single, narrower track (Fig. 4F - 4H). This indicates that Ppa-SYP-1 is a transverse filament protein arranged in a head-to-head orientation, spanning the width of the SC. Ppa-SYP-1 appears to be a functional homolog of *C. elegans* SYP-1: they are both transverse filament proteins that contain similar coiled-coil domain structures, predicted to be 55 and 51 nm long, respectively (Fig. 4I, compare to Fig. 2A, SYP-1; (14, 43)), and both harbor Polo-Box Domains in their C-termini (STP at position 451 - 453 for *C. elegans* SYP-1 (44) and positions 658 – 660 and 681 – 683 for Ppa-SYP-1). This is despite the fact that Ppa-SYP-1 has no significant homology to *C. elegans* SYP-1, or, for that matter, to any known SC protein.

## Discussion

The discrepancy between the ultrastructural and functional conservation of the SC on one hand, and the sequence divergence of its constituent proteins on the other, have long baffled chromosome biologists. Here we show that while analysis of per-site identity and similarity reveals high degree of sequence divergence, SC proteins exhibit significant and widespread conservation of protein length and of coiled-coil domains. SC proteins thus provide a clear example of a first-order amino acid fitness landscape tolerant of mutations accompanied by second-order fitness landscapes (properties derived from primary amino acid sequence) that impose strong evolutionary constraints. This provides an alternative framework to understand rapid divergence of proteins performing essential functions, in addition to the more well-documented processes of adaptive evolution and positive selection (45).

In the SC, the strong selection acting on coiled-coils likely reflects biophysical and structural properties that are conserved across phyla. Coiled-coils in transverse filament proteins help determine the 100nm space between the parental chromosomes (13, 15). However, this role by itself cannot explain the prevalence and conservation of coiled-coils in SC proteins that are not transverse filaments. Moreover, it cannot explain conserved disruptions in the coiled-coils (Fig. 2). An attractive hypothesis is that the specific arrangement of the coiled-coils and their disruptions determines other aspects of the 3-dimensional architecture of the SC lattice (31), such as its axial depth or the uniform lateral spacing between the ladder rungs (Fig. 1A). In that case, many functions of the SC could be maintained by swapping endogenous coiled-coils with orthologous or synthetic ones of similar length and arrangement.

An alternative, non-mutually exclusive role for the coiled-coils is promoting phase-separation. Despite its ordered appearance, the SC in worms, flies and yeast has recently been shown to assemble through phase-separation (also referred to as condensation; (46)). Constituent subunits of condensates, including SC proteins, can enter and exit condensates and move within them. Coiled-coils can facilitate phase-separation (47, 48), potentially by promoting multivalent interactions (49, 50). This is consistent with the poor per-site conservation of SC proteins, since multivalent interactions can rely on molecular features exhibited by groups of amino acids (e.g., charge or hydrophobicity) rather than tight, ‘lock-and-key’ interfaces formed by specific tertiary structures. Tellingly, SPD-5, which shares an evolutionary signature with SC proteins in *Caenorhabditis* (Fig. S5), has been shown to promote microtubule nucleation in mitosis through condensation (51). Given the growing number of characterized condensates in the cell (52) it is tempting to speculate that many of their constituent proteins are subject to unconventional evolutionary pressures that are not apparent in their primary amino acid sequence. Prime candidates to examine in this regard are disordered protein domains (Fig. S4) which can drive phase-separation (53, 54), and tend to evolve more rapidly than their ordered counterparts (55).

Despite rapidly evolving amino acid sequence, purifying selection acts to limit length variation in SC proteins (Fig. 2 and 3). The indels that are present in SC proteins are depleted within the coiled-coils (Fig. 2D, Table S2). Such uncoupling is unusual, since indels and substitutions typically occur together (56, 57). In fact, selection acting on indels has been demonstrated only in a handful of cases. One such case are sperm ion channels in primates, rodents and flies, where positive selection for indels yielded N-terminal tails of highly varied lengths (58–60). Robust genome-wide identification of indels is complicated by assembly and annotation errors that can be mistaken for indels (see Methods). Future work to develop methods to test for selection against indels is likely to shed light on these evolutionary dynamics of protein length variation, and on the mechanisms underlying them.

Our ability to detect the unconventional conservation of SC proteins relied on the ultrastructural conservation of the SC across eukaryotes. This knowledge was gained through the widespread application of electron microscopy, and the serendipitous ability to observe the SC without any molecular knowledge of its constituent subunits. Unlike the SC, however, much of our current understanding of cellular organization relies on the application of molecular tools (e.g., antibodies, tagged transgenes). These efforts are often informed and directed by conservation of primary amino acid sequence to select ‘interesting’ targets for cell biological and genetic experiments, and to actively avoid so-called orphan genes. Our work shows that by focusing our explorations under the streetlamp that are BLAST searches we might be ignoring conserved cellular structures and consequential biological processes.

## Materials and Methods

### Identification of SC proteins in Caenorhabditis

To identify SC proteins in *Caenorhabditis* we used *C. elegans* SYP-1 – SYP-5 as queries in BLASTP and tBLASTn searches of 19 species in the *Elegans* group and *Drosophilae* supergroup. For the remaining six species, which are more distantly related to *C. elegans*, we used SYP-1 – SYP-5 sequences from all *Elegans* and *Drosophilae* supergroups species as BLAST queries. We compared the syntenic location (5’ and 3’ neighbor genes) of each BLAST hit and built gene-specific phylogenies to confirm orthology (Fig. S2, supplementary data). In several cases (23/125, 18%), gene annotations were either absent or incorrect. For example, three annotations merged two genes and seven had obvious errors in intron/exon boundaries. Uncorrected, these annotations would have manifested as apparent indels. We corrected these errors manually using expression data when available and by alignment to closely related species (supplementary data). In a few cases, ambiguities in genome assemblies prevented us from generating confident gene models. *C. japonica, C. inopinata, C. virilis, C. angaria and C. monodelphis* SYP-4 all share significant homology to other SYP-4s. However, their gene models reside on short scaffolds, the edges of scaffolds or contain ambiguous bases. Similarly, *C. tropicalis, C. waitukubuli, C. japonica* and *C. angaria* SYP-5 all contained ambiguities. We scored each of these genes as present, but did not use them in further analyses. Alignments used for gene-specific phylogenies were generated using ClustalW (61) implemented in Geneious Prime (version 2019.0.4). Maximum likelihood phylogenies were generated with PhyML (version 3.3.20200621) with the LG amino acid substitution model and 100 replicates for bootstrap support (62).

### Testing for positive selection

We selected the Elegans supergroup as an appropriate subset of species to test for recurrent positive selection (63). We used ClustalW as described above to make alignments of each SC protein and of SMC-1/3 (controls) from 12 Elegans group species (*C. zanzibari, C. tribulationis, C. sinica, C. briggsae, C. nigoni, C. remani, C. latens, C. doughertyi, C. brenneri, C. tropicalis, C. inopinta,* and *C. elegans).* Since *C. elegans* contains a SYP-5 paralog, SYP-6, we excluded *C. elegans* SYP-5/6 from the SYP-5 protein alignment. We generated corresponding nucleotide alignments using Pal2Nal (64). Each alignment, along with an *Elegans* group species tree (65) was used as input to the CodeML sites model of PAML (28) (supplementary data). We compared models M1 (neutral) and M2 (selection), M7 (dN/dS < 1) and M8 (dN/dS < 1, plus an additional category of dN/dS > 1), and M8a (dN/dS ≤ 1) and M8. We tested for significance in each comparison using a likelihood ratio test. We ensured that our results were robust to codon substitution model and starting dN/dS by running each test with two codon models (F3×4 and the codon table derived from each alignment) and with two starting dN/dS values (dN/dS = 0.4 and dN/dS = 1.5). To ensure that our lack of detection of positive selection was not due to the relatively high divergence of the *Elegans* group species, we repeated the analysis excluding *C. elegans* and *C. inopinata,* the most divergent members of the Elegans species group. We found no evidence for positive selection using this less-diverged species set (Table S1). We also tested for pervasive positive selection using a Fixed Effects Likelihood method (66) implemented at datamonkey.org. We used the same nucleotide alignments that we used for PAML analysis of the full *Elegans* group as input. Alignment-wide average pairwise percent identity for SYP-1 – SYP-5 and SMC-1/3 (Fig. 1C) and percent identity by site (Fig. 2A) was calculated in Geneious. Sliding window percent identity (window size = 10 amino acids) was calculated in R (version 4.0.2).

### Generating orthogroups, making alignments and calculating divergence

We used OrthoFinder (67, 68) with default parameters to create groups of orthologous proteins (orthogroups) from *Drosophila, Eutherian* mammalian, and *Pristioncus* genomes or proteomes (Table S3). *Caenorhabditis* orthogroups were generated previously (65). We removed paralogous proteins from each orthogroup by removing the species containing the duplicate gene from that orthogroup. For *Caenorhabditis, Drosophila* and mammalian analyses, we only analyzed orthogroups that contained proteins from at least half of the possible species after removing paralogs (13, 15 and 7 species, respectively). In an effort to aid identification of SC proteins in *Pristionchus,* and since there are only eight *Pristionchus* genomes available, we did not apply these filtering steps to *Pristionchus* orthogroups. This resulted in 9924 *Caenorhabditis* orthogroups, 11622 *Drosophila* orthogroups, 18470 mammalian orthogroups and 28042 *Pristionchus* orthogroups. We aligned all orthogroups using ClustalW implemented in MEGA (69, 70). We also used MEGA to estimate overall mean amino acid substitutions per site, calculated for all pairwise combinations, under a Poisson substitution model for each aligned orthogroup. We assumed rate variation among sites followed a Gamma distribution with Gamma parameter = 2.00.

### Coiled-coil domain prediction and coiled-coil conservation scores

We used Paircoil2 with window size = 28 for all coiled-coil domain predictions (30). To calculate the coiled-coil conservation score, we aligned the coiled-coil scores (1 – Paircoil2 score) for each orthogroup based on the amino acid alignment using a custom Python script. Alignment columns with fewer than 80% of species represented were removed in the plots shown throughout. For *Pristionchus* analyses, we removed alignment columns that had fewer than seven out of eight species represented. For proteins of interest (SYP-1 – SYP-5, C(3)G, Corolla, Corona, SYCP1, SYCE1, SYCE2, SYCE3, TEX12, SIX60S1 and all candidate *Pristionchus* SC proteins), aligned coiled-coil scores were visualized using R. Average and standard deviation of aligned coiled-coil scores was calculated and visualized in R. We then took the minimum value at each alignment position and averaged it across the length of the alignment. We refer to this averaged value the coiled-coil conservation score. Conserved coiled-coils (grey ovals in Fig. 2A, Fig. 3, Fig. S6 and S7) were defined as regions of the coiled-coil plots at least 21 amino acids long where the average coiled coil score was > 0.8 and the standard deviation of the coiled-coil score was < 0.1.

### Coefficient of variation of protein length

Coefficient of variation of protein length was calculated as the standard deviation of protein length in each orthogroup divided by the mean length of the proteins in that group.

### Statistical tests comparing SC proteins to the proteome

We used a Wilcoxon rank sum test to compare median amino acid substitutions per site (Fig. 1C), coiled-coil conservation score (Fig. 2B) and coefficient of variation of protein length (Fig. 2C) of SC proteins to the rest of the proteome.

### Disordered domain prediction

We used PONDR VL3 for all disordered domain predictions (71). We aligned PONDR VL3 scores for SYP-1 – SYP-5 using a custom Python script and visualized the aligned scores using R. Average and standard deviation of the aligned PONDR VL3 scores were calculated and visualized in R.

### Indel analysis

We used ClustalW to generate an alignment of each SC protein (SYP-1 – SYP-5) in the *Elegans* supergroup (supplementary data). We constructed a neighbor-joining tree corresponding to each alignment. We then scanned each alignment and manually counted instances of insertions and deletions along the phylogeny. Each position in the alignment was then defined as either indel-containing or indel-lacking. Separately, each alignment position was classified as either part of or not part of a coiled-coil domain based on the coiled-coil score at each position in *C. elegans.* For the SYP-5 alignment, we excluded *C. elegans* SYP-5 because *C. elegans* contains a SYP-5 paralog, SYP-6. Therefore, we defined the coiled-coil alignment columns based on *C. briggsae* SYP-5. For statistical analysis, we generated a 2 × 2 contingency table comparing the prevalence of indel-containing positions to the coiled-coil domains and calculated a two-sided p-values from Fisher’s test from combined data from SYP-1 – SYP-5. Our null hypothesis was that indels would be equally likely to occur in coiled-coil and non-coiled-coil domains.

### Hierarchical clustering

Proteins were assigned a point in a 3-dimensional Cartesian coordinate system where × = amino acid substitutions per site, y = coiled-coil conservation score and z = coefficient of variation of protein length. We used the dist function in the stats package in R to generate a dissimilarity matrix based on Euclidian distance between each point. The dissimilarity matrix was used to perform complete linkage clustering using hclust, also in R. We plotted the results as a dendrogram with 15 leaves (Fig. S5).

### Enrichment analysis

We used the Database for Annotation, Visualization and Integrated Discovery (DAVID v6.8) to test for enrichment of Gene Ontology (Biological Processes sub-ontology) in the two dendrogram leaves containing SC proteins (Fig. S5A, blue and green dots). Significantly enriched categories were: embryo development ending in birth or egg hatching (GO:0009792, p-value = 9.5 × 10^−9^), synaptonemal complex assembly (GO:0007130, p-value = 0.0035), meiotic nuclear division (GO:0007126, p-value = 0.0046), meiotic chromosome segregation (GO:0045132, p-value = 0.0074), and mitotic spindle organization (GO:0007052, p-value = 0.0079). P-values reported were Benjamini corrected.

### *P. pacificus* maintenance

*P. pacificus* strains were grown at 20°C on NGM agar with *E. coli* OP50. *P. pacificus* strain PS312 (obtained from the CGC) was used for injections and as a wild-type control. All strains were homozygous except for the *Ppa-syp-1* knockout which was maintained as heterozygote. In order to perform progeny counts and DAPI body counts of homozygous *Ppa-syp-1* knockout animals (experiments described in more detail below), we singled animals to identify heterozygotes, and identified the homozygous knockout animals among their progeny based on the presence of laid eggs but no viable progeny (compared to their wild-type or heterozygous siblings which had many viable progeny). These homogyzous knockout animals were used for subsequent experiments and genotyped by PCR.

### Prioritizing SC candidates for knockout in *P. pacificus*

We prioritized our list of *Pristionchus* candidate SC proteins using the following criteria: 1) The gene must be single copy in *P. pacificus.* 2) The protein must not have a significant BLAST match to any protein in *C. elegans.* Since *C. elegans* SC proteins have no significant homology to any *P. pacificus* protein, we reasoned that the reverse would also be true. 3) We prioritized candidates that showed gonad-specific enrichment based on RNA tomography (72). However, since the RNA tomography study (72) used gene names from previous strand-specific transcriptome assemblies (73, 74) and our analysis was based on the El Paco V2 genome assembly, we first identified candidates whose expression was enriched in J4 *versus* J1 *P. pacificus* using RNAseq data (NCBI SRA: PRJNA628502, Michael Werner, personal communication). This left 68 adult enriched candidates. We used each of these 68 candidates as queries in a BLAST search against the strand-specific transcriptome to identify a gene name that corresponds to the RNA-tomography data, and selected genes that were gonad enriched. We were left with eight candidates, including Ppa-SYP-4 (42).

### Construction of *P. pacificus* strains

*P. pacificus* J4 animals were grown for 24 hours at 20°C prior to injection. Injection mix was prepared as follows: to make the RNA mix, we combined 4 μl tracrRNA (IDT, 200 μM) and 4 μl crRNA (200 μM) and incubated at 95°C for 5 minutes. We let the RNA mix cool on the benchtop for 5 minutes. To make the injection mix, we combined 3.5 μl RNA mix, 1 μl Cas9 protein (IDT, 10 μg/μl), 3 μl single-stranded DNA repair template (200 μM) and 1 μM DNA duplex buffer (IDT). To detect CRISPR events, we screened the pooled F1 progeny of injected P_0_ animals for the loss of an endogenous restriction site near the Cas9 cut site (so-called “jackpot” plates). We then singled F1s from jackpot plates for subsequent genetic analysis.

Construction of null mutants in candidates was done by inserting a premature stop codon in the N-terminus of each candidate. Stop codons were generated either via homology directed repair or by random indels generated by Cas9. *Ppa-syp-1* had a stop codon 94 bp from the N-terminal ‘ATG’, *PPA10754* had a 10 bp insertion 6 bp from the N-terminus resulting in a frame shift and a stop codon after 75 bp, and *PPA35551* had a N-terminal deletion of 222 bp resulting in a stop codon after 78 bp. Construction of Ppa-SYP-1 strains tagged with OLLAS and HA was done by inserting each tag within the *Ppa-syp-1* coding sequence using homology directed repair. The HA tag was inserted 5’ of the extended coiled-coil and is flanked by three glycine linkers on each side. The OLLAS tag was inserted 3’ of the extended coiled-coil. Both tagged strains are homozygous viable and produce similar numbers of viable self-progeny to wild-type *P. pacificus* (Fig. 4D). All edits were verified by Sanger sequencing. See supplementary data for a list of gene-specific crRNAs, primers and restriction enzymes used for genotyping.

### Progeny counts

We picked at least 10 single *P. pacificus* J4 animals onto individual plates. We transferred each animal to a new plate every day for five days. The resulting progeny on each plate were counted as adults four days after the parent was moved off of the plate. Progeny counts were performed at 20°C.

### Imaging

We dissected age-matched *P. pacificus* hermaphrodites (24hr post-J4) in 30 μl 1x Egg Buffer, essentially as described in (75) with 0.01% Tween-20 and 0.005% tetramisole on a 22 × 22 mm coverslip. To fix, we added equal volume of a 2% formaldehyde solution in 1× Egg Buffer and incubated for one minute. We removed nearly all dissection/fixation solution from the sample and picked up the coverslip with a HistoBond microscope slide (VWR). Samples were then frozen on dry ice. After freezing, we snapped the coverslip off and immediately immersed samples in −20°C methanol for 1 minute. We then washed samples 3 × 5 minutes in PBST (0.1% Tween-20) and incubated in primary antibodies overnight at 4°C. We washed samples 3 × 5 minutes in PBST and incubated in secondary antibodies for 2 hours at room temperature. Samples were washed for 10 min in PBST and for 10 min in DAPI (5 μg/μl). Samples were mounted in NPG-glycerol. Antibodies were used as follows: (primaries) mouse anti-HA (Roche 12CA5) 1:500, rat anti-OLLAS clone L2 (Invitrogen) 1:200, (secondary for confocal) donkey anti-mouse Alexa 488 (Jackson ImmunoResearch) 1:500, (secondaries for STED) goat anti-mouse STAR RED (Aberrior) 1:100, donkey anti-rat Alexa 594 (Jackson ImmunoResearch 1:500). For STED microscopy, samples were prepared as above except we omitted DAPI staining and mounted in Aberrior Mount liquid antifade (Aberrior) instead of NPG-glycerol. STED images were acquired with an Aberrior STEDYCON mounted on a Nikon Eclipse Ti microscope with a 100× 1.45 NA oil objective. Confocal images were acquired on a Zeiss LSM880 with Airyscan and a 63× 1.4 NA oil objective. STED images are a single z-section and confocal images are partial maximum intensity projections.

### DAPI body counting

48hr post-J4 *P. pacificus* hermaphrodites were dissected and stained as described above, except for omitting antibody staining. Oocytes with condensed chromosomes (typically in the −1 or −2 position) were imaged with a confocal z-stack. DAPI bodies were counted from 3D renderings in Zen Blue.

### Line scan measurements

Pixel intensities from STED images of *Ppa-syp-1:OLLAS* (n = 13 chromosomes) and *HA::Ppa-syp-1* (n = 10 chromosomes) were measured via line scan perpendicularly across the SC in ImageJ (version 2.1.0/1.53c). Pixel intensities were normalized to the maximum value in each line scan. Line scans from multiple chromosomes were aligned using the center of the SC as a reference. Line scan averages were calculated in R.

## Supporting information

Supplemental data - alignments for indel analysis

Supplemental data - alignments for maximum likelihood phylogenies

Supplemental data - alignments for PAML

Supplemental data - CRISPR reagents

Supplemental data - manually curated genes

Supplemental data - proteomes

Supplemental data - synteny

Supplemental data - SYP protein sequences

Supplemental data - trees for indel analysis

## Authors contributions

LEK performed all evolutionary and bioinformatic analysis. LEK and HDC performed *P. pacificus* experiments. LEK and OR conceptualized the project and wrote the paper.

## Acknowledgments

We would like to thank Michael Werner for *P. pacificus* advice, reagents and RNAseq data, Lewis Stevens for *Caenorhabditis* orthogroups, The University of Utah Center for High Performance Computing for computational resources, and the Rog lab, Harmit Malik lab, Sophie Caron, Jon Seger and Fred Adler for discussions in various stages of this project. We would also like to thank Yuval Mazor, Nitin Phadnis, Talia Karasov and Michael Werner for critical reading of the manuscript, and Sara Nakielny for comments on the manuscript and editorial work. Worm strains were provided by the CGC, which is funded by NIH Office of Research Infrastructure Programs (P40 OD010440). LEK is supported by the Developmental Biology Training Grant from NICHD (T32HD007491). Work in the Rog lab is supported by R35GM128804 grant from NIGMS, and start-up funds from the University of Utah.

## Supplementary Figures and Tables

**Fig. S1:**
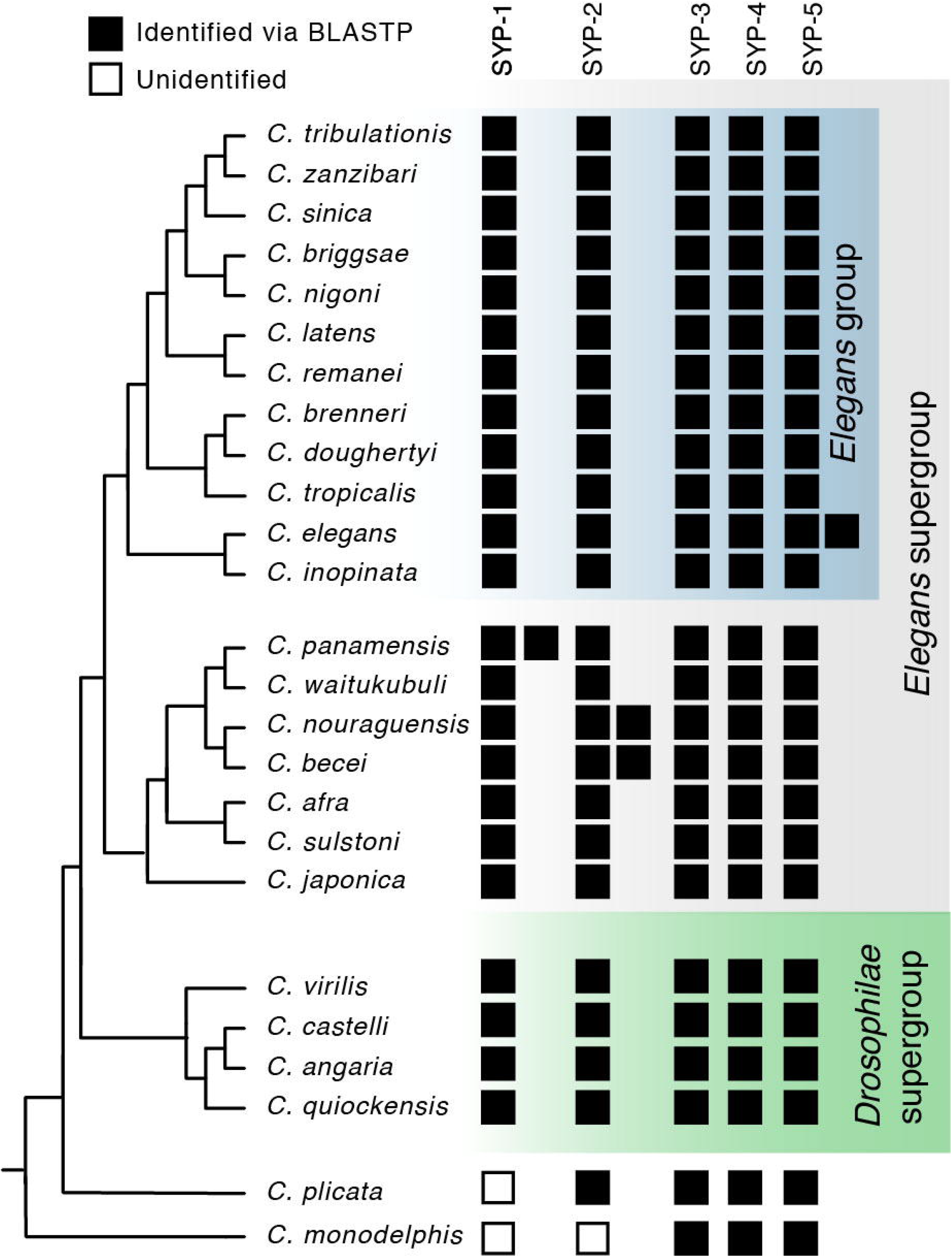
Identification of *Caenorhabditis* SC proteins. *Caenorhabditis* species tree including all species investigated in this study. Presence of SC proteins is listed to the right of each species. Filled box = present, unfilled box = no ortholog could be identified. Related to Fig. 1B.

**Fig. S2:**
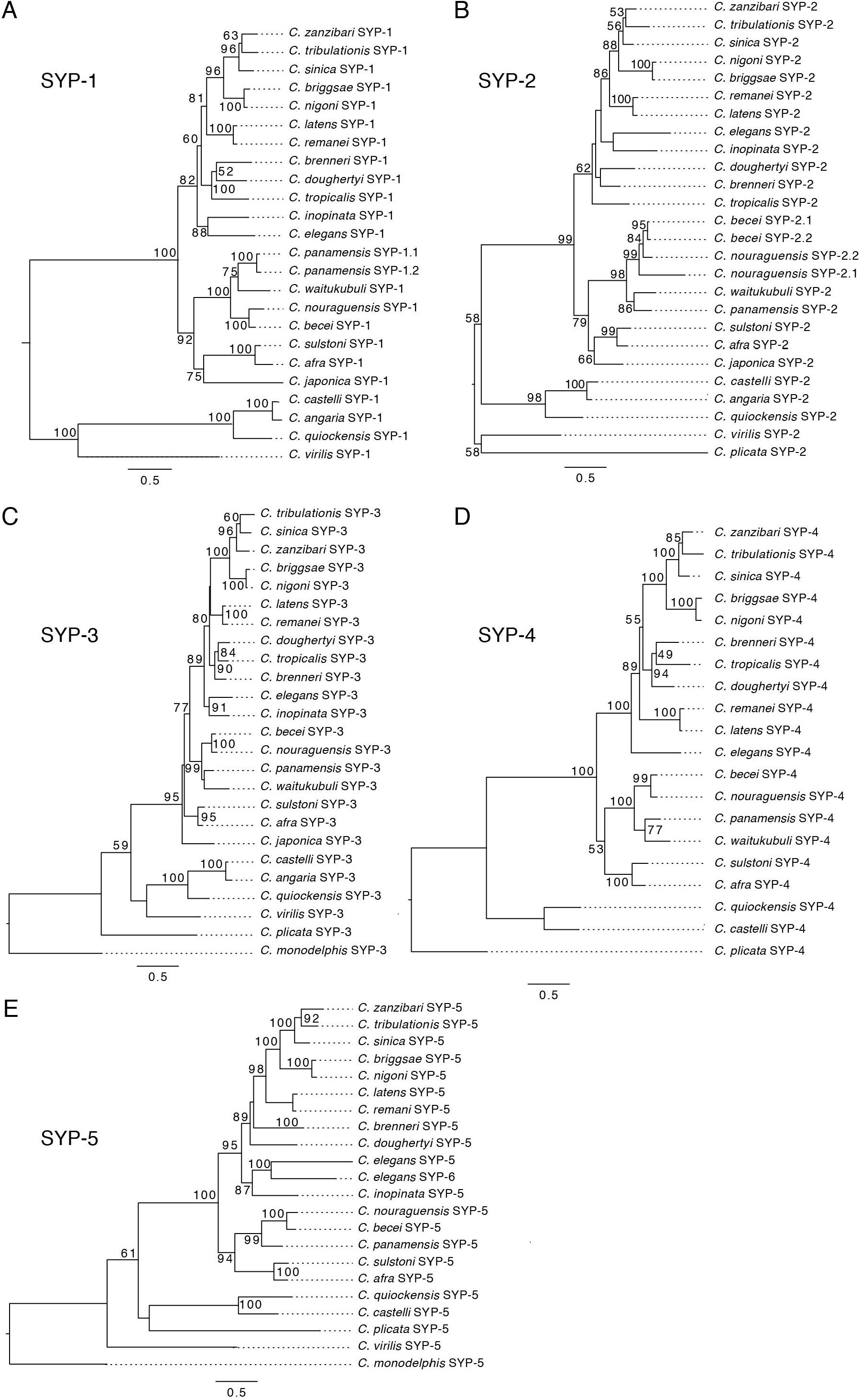
Maximum likelihood phylogenies of *Caenorhabditis* SC proteins. Maximum likelihood phylogenies of SYP-1 (A) – SYP-5 (E). Each phylogeny is rooted on the common ancestor of *Caenorhabditis.* Bootstrap values above 50 are displayed at each node. Scale bar represents amino acid substitutions per site.

**Fig. S3:**
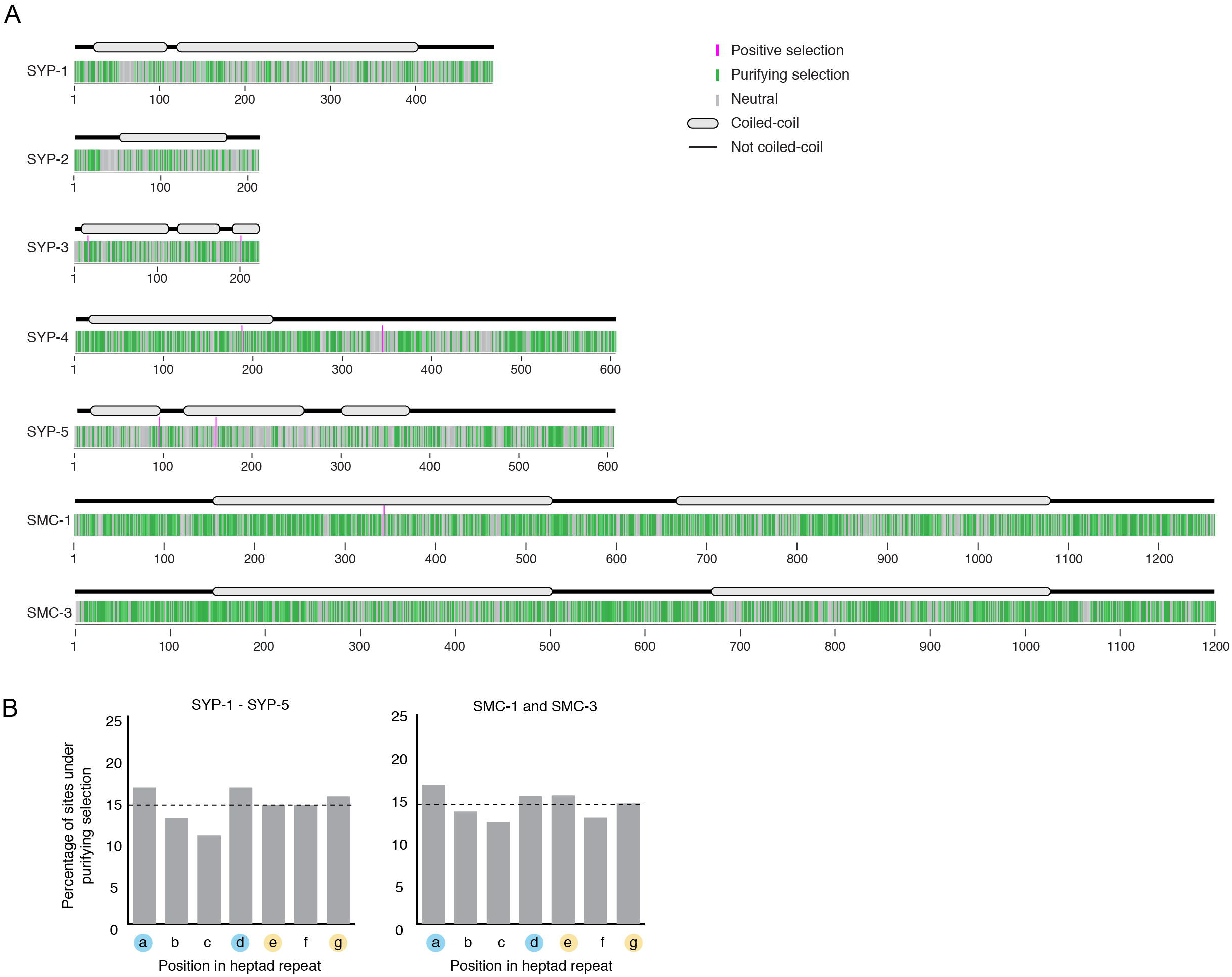
Map of sites of under purifying, neutral or positive selection. (A) Map of sites under positive (pink), purifying (green) or neutral selection (grey) identified using Fixed Effects Likelihood model from HyPhy for SYP-1 through SYP-5 and SMC-1/3. Gene models showing conserved coiled-coils (grey ovals) are shown above. (B) Graph of percentage of sites within coiled-coil domains evolving under purifying selection. Consistent with the requirements for coiled-coils - hydrophobic amino acids in positions one and four (designated a and d, blue circles) and charged or polar amino acids in positions five and seven (designated e and g, yellow circles) - we found an enrichment of sites under purifying selection in the first, fourth and seventh (a, d and g) position of SYP-1 through SYP-5, and in the first, fourth and fifth (a, d and e) position of SMC-1/3. However, these differences were not statistically significant (SYP-1 – SYP-5 p-value = 0.12, SMC-1/3 p-value = 0.15). Horizontal dashed line indicates the expected percentage if sites under purifying selection were randomly distributed.

**Fig. S4:**
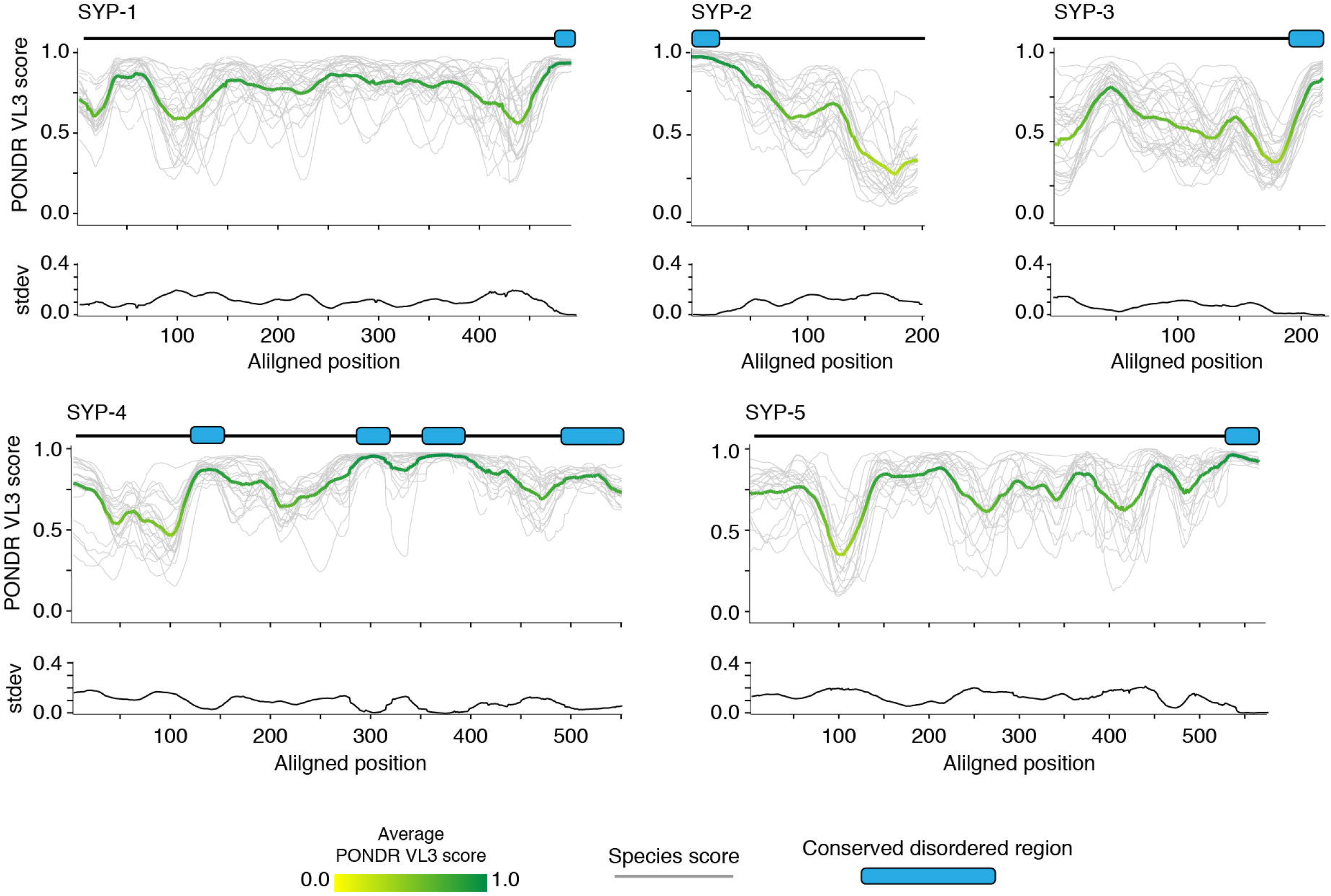
SC proteins have limited regions of conserved disorder. PONDR VL3 score (top) and standard deviation of VL3 (bottom) at each aligned position for SYP-1 – SYP-5. Green and yellow line = average score at each position. Grey lines = scores for each species. Gene models depicting conserved disordered domains (blue-filled ovals) are shown above each PONDR VL3 plot.

**Fig. S5:**
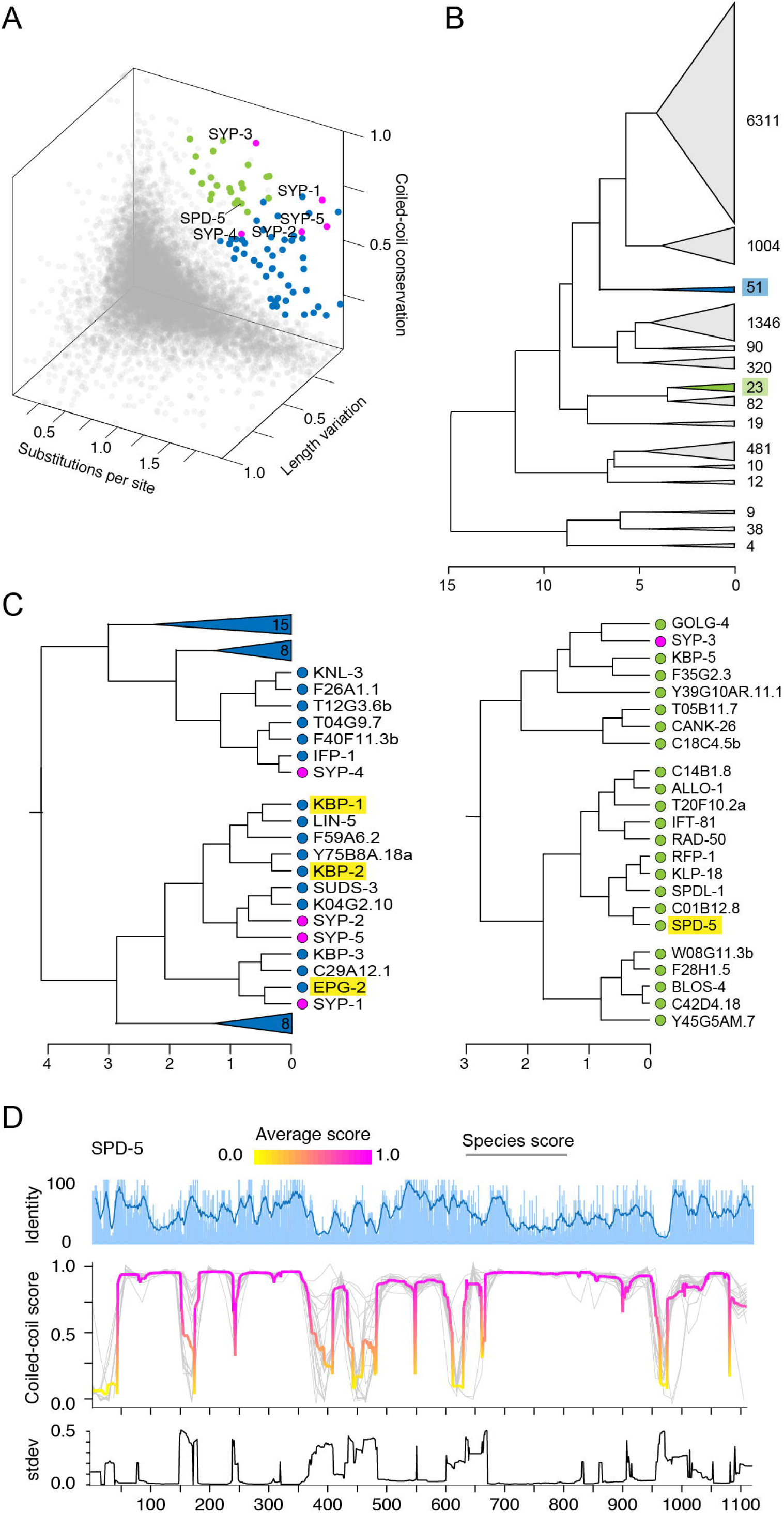
SC proteins have an unconventional evolutionary signature. (A) 3D scatter plot from Fig. 3A re-printed with proteins that cluster with SC proteins based on hierarchical clustering highlighted in blue and green. (B) Hierarchical clustering of all *Caenorhabditis* proteins based on amino acid substitutions per site, coiled-coil conservation score and coefficient of variation of protein length. SC proteins cluster outside of the main clusters of 6311 and 1004 proteins. Blue and green highlighted clusters contain SC proteins. (C) Partially expanded clusters containing SC proteins from (B), similarly colored in blue and green. Gene names are listed at the tip of each branch. Enrichment analysis of the clusters containing SC proteins identified enrichment of proteins involved in mitotic spindle organization (highlighted in yellow, p-value = 0.0079), including SPD-5. (D) Sliding window percent identity and coiled-coil plot for SPD-5. Grey lines = individual species score, Magenta and yellow line = average score at each position. Note conservation of the coiled-coils and disruptions in them, despite overall low sequence identity.

**Fig. S6:**
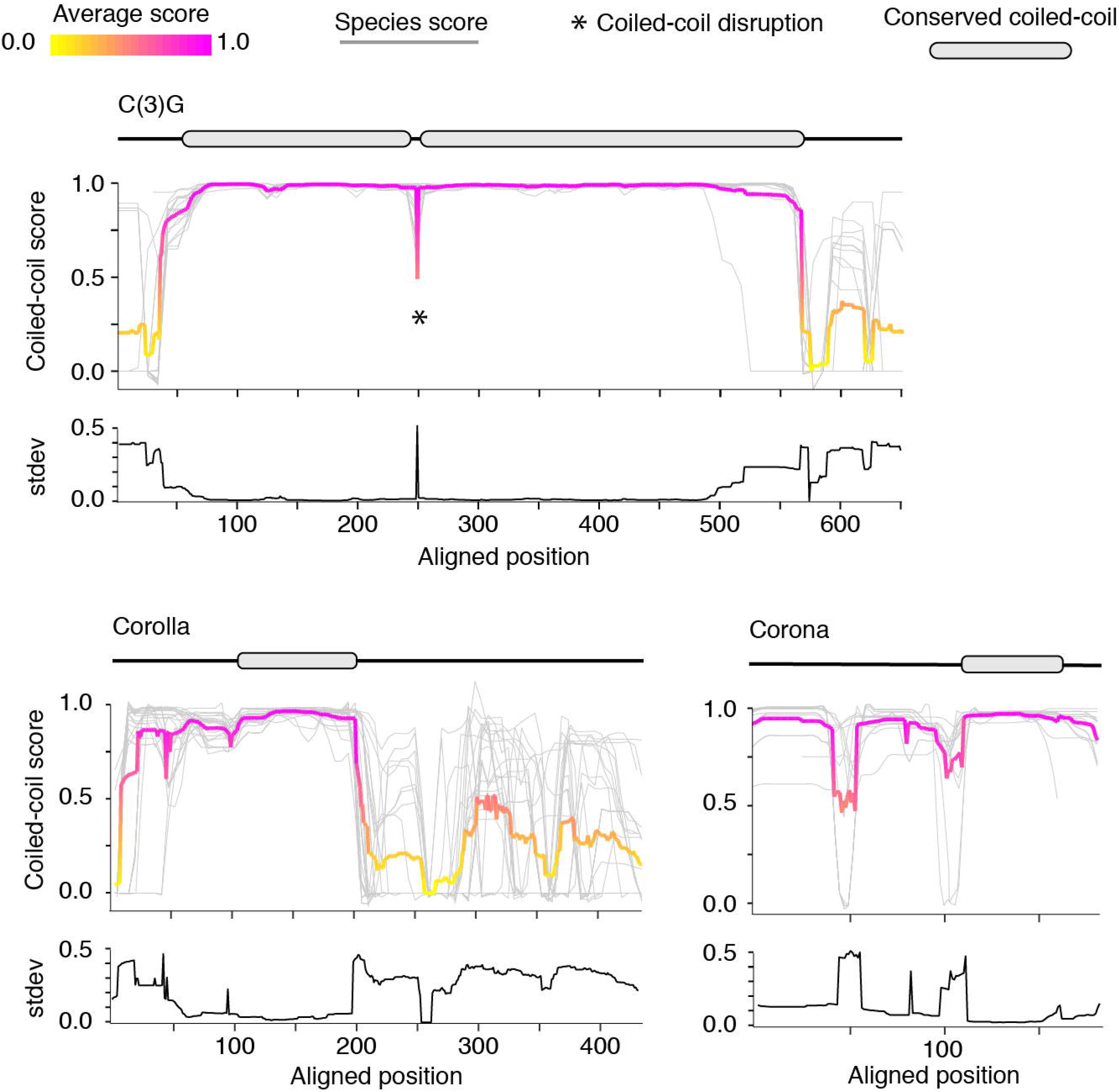
Coiled-coil plots for *Drosophila* SC proteins. Coiled-coil plots for C(3)G, Corolla and Corona used to identify conserved coiled-coil domains (grey ovals) displayed in Fig. 3B. Grey lines = individual species score, Magenta and yellow line = average score at each position. Standard deviation of coiled-coil score is shown below each coiled-coil plot.

**Fig. S7:**
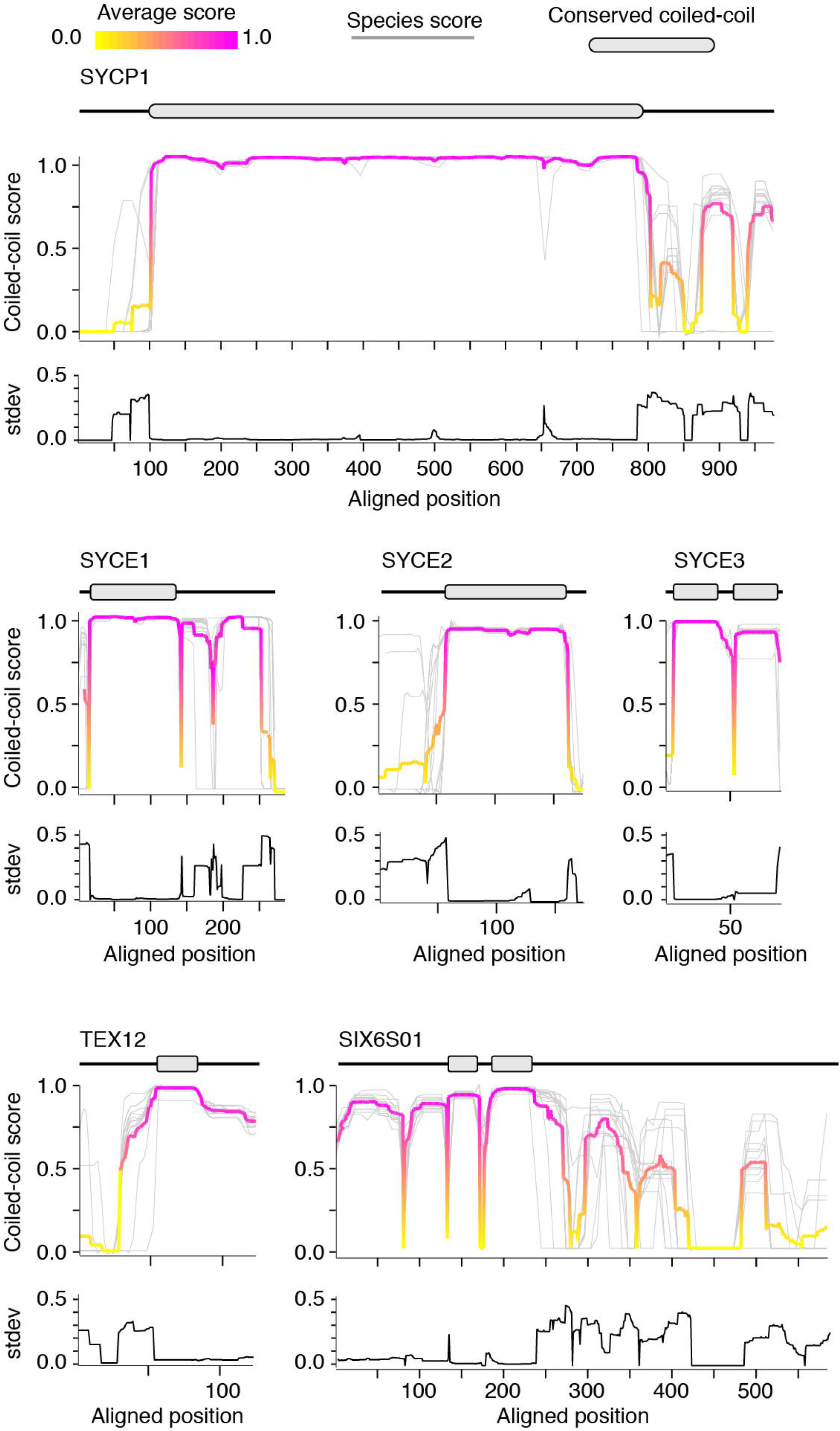
Coiled-coil plots for mammalian SC proteins. Coiled-coil plots for SYCP1, SYCE1 – 3, TEX12, and SIX6S01 used to identify conserved coiled-coil domains (grey ovals) displayed in Fig. 3C. Grey lines = individual species score, Magenta and yellow line = average score at each position. Standard deviation of coiled-coil score is shown below each coiled-coil plot.

**Table S1.** Summary table of tests for positive selection on SC proteins and SMC-1 and SMC-3 (controls) in the Elegans group species of *Caenorhabditis* (Fig. S1). P-values from likelihood ratio tests comparing CodeML models M1 vs M2, M7 vs M8, and M8a vs M8 are listed. Each comparison tests the fit of the data to a model that does not allow positive selection (M1, M7 and M8a) to a model that does allow positive selection (M2 and M8). P-values from likelihood ratio tests using a smaller set of species are also provided for SYP-1 – SYP-5 (see methods). The number of sites under positive or negative selection in each protein according to the Fixed Effects Likelihood analysis from HyPhy with p-value < 0.05 are also displayed.

|  | PAML<br>CodeML |  |  |  | HyPhy<br>Fixel Effects Likelihood |  |
| --- | --- | --- | --- | --- | --- | --- |
| Gene | Number of<br>species | M1 vs M2<br>p-value | M7 vs M8<br>p-value | M8a vs M8<br>p-value | sites under<br>pos. selection | sites under<br>neg. selection |
| SYP-1 | 12 | 1.00 | 0.99 | 0.60 | 0/451 | 177/451 (39%) |
| SYP-2 | 12 | 1.00 | 0.90 | 0.56 | 0/203 | 75/202 (37%) |
| SYP-3 | 12 | 0.97 | 0.54 | 0.54 | 2/213 | 109/213 (51%) |
| SYP-4 | 11 | 1.00 | 0.26 | 0.48 | 2/550 | 277/550 (50%) |
| SYP-5 | 10 | 1.00 | 0.99 | 0.59 | 2/554 | 173/554 (31%) |
| SMC-1 | 11 | 1.00 | 0.85 | 0.54 | 1/1300 | 805/1300 (62%) |
| SMC-3 | 12 | 1.00 | 0.51 | 0.41 | 0/1243 | 760/1243 (61%) |
| SYP-1 | 10 | 1.00 | 1.00 | 0.67 | - | - |
| SYP-2 | 10 | 1.00 | 1.00 | 0.77 | - | - |
| SYP-3 | 10 | 1.00 | 1.00 | 0.81 | - | - |
| SYP-4 | 10 | 1.00 | 0.14 | 0.64 | - | - |
| SYP-5 | 9 | 1.00 | 1.00 | 0.56 | - | - |

**Table S2.** Contingency table showing the number of alignment positions containing indels and lacking indels inside versus outside the coiled-coil domain of each SC protein. Two tailed p-value from Fishers exact test is shown in the last column. Total number of insertions and deletions are depleted in the coiled-coil domains of SYP-1, SYP-4 and SYP-5.

|  | Inside coiled-coil |  | Outside coiled-coil |  |  |
| --- | --- | --- | --- | --- | --- |
| Protein | Aln. positions<br>with indels | Total aln.<br>positions | Aln. positions<br>with indels | Total aln.<br>positions | p-value |
| SYP-1 | 30 | 407 | 54 | 157 | < 0.0001 |
| SYP-2 | 4 | 104 | 3 | 92 | 1.0 |
| SYP-3 | 8 | 251 | 0 | 33 | 0.6 |
| SYP-4 | 3 | 237 | 31 | 161 | < 0.0001 |
| SYP-5 | 14 | 372 | 14 | 131 | 0.0063 |
| Total | 59 | 1371 | 102 | 574 | < 0.0001 |

## References

1. S. L. Sawyer, L. I. Wu, M. Emerman, H. S. Malik, Positive selection of primate TRIM5alpha identifies a critical species-specific retroviral restriction domain. Proc. Natl. Acad. Sci. U. S. A. 102, 2832–2837 (2005).

2. P. S. Mitchell, et al., Evolution-guided identification of antiviral specificity determinants in the broadly acting interferon-induced innate immunity factor MxA. Cell Host Microbe 12, 598–604 (2012).

3. M. D. Daugherty, H. S. Malik, Rules of engagement: molecular insights from host-virus arms races. Annu. Rev. Genet. 46, 677–700 (2012).

4. K. A. Lewis, D. S. Wuttke, Telomerase and telomere-associated proteins: structural insights into mechanism and evolution. Structure 20, 28–39 (2012).

5. F. Wang, et al., The POT1–TPP1 telomere complex is a telomerase processivity factor. Nature 445, 506–510 (2007).

6. M. J. Moses, Chromosomal structures in crayfish spermatocytes. J. Biophys. Biochem. Cytol. 2, 215–218 (1956).

7. D. W. Fawcett, The fine structure of chromosomes in the meiotic prophase of vertebrate spermatocytes. J. Biophys. Biochem. Cytol. 2, 403–406 (1956).

8. C. B. Gillies, P. B. Moens, The synaptonemal complex in higher plants. CRC Crit. Rev. Plant Sci. 2, 81–116 (1984).

9. A. T. Carpenter, Electron microscopy of meiosis in Drosophila melanogaster females. I. Structure, arrangement, and temporal change of the synaptonemal complex in wild-type. Chromosoma 51, 157–182 (1975).

10. A. J. MacQueen, M. P. Colaiácovo, K. McDonald, A. M. Villeneuve, Synapsis-dependent and -independent mechanisms stabilize homolog pairing during meiotic prophase in C. elegans. Genes Dev. 16, 2428–2442 (2002).

11. K. Schmekel, J. Wahrman, U. Skoglund, B. Daneholt, The central region of the synaptonemal complex in Blaps cribrosa studied by electron microscope tomography. Chromosoma 102, 669–681 (1993).

12. M. J. Moses, L. B. Russel, N. L. Cacheiro, Mouse chromosome translocations: visualization and analysis by electron microscopy of the synaptonemal complex. Science 196, 892–894 (1977).

13. M. Sym, G. S. Roeder, Zip1-induced changes in synaptonemal complex structure and polycomplex assembly. J. Cell Biol. 128, 455–466 (1995).

14. K. S. Tung, G. S. Roeder, Meiotic chromosome morphology and behavior in zip1 mutants of Saccharomyces cerevisiae. Genetics 149, 817–832 (1998).

15. K. K. Billmyre, et al., X chromosome and autosomal recombination are differentially sensitive to disruptions in SC maintenance. Proc. Natl. Acad. Sci. U. S. A. 116, 21641–21650 (2019).

16. R. Ollinger, M. Alsheimer, R. Benavente, Mammalian protein SCP1 forms synaptonemal complex-like structures in the absence of meiotic chromosomes. Mol. Biol. Cell 16, 212–217 (2005).

17. J. B. Woodruff, Assembly of Mitotic Structures through Phase Separation. J. Mol. Biol. 430, 4762–4772 (2018).

18. M. E. Hurlock, et al., Identification of novel synaptonemal complex components in C. elegans. J. Cell Biol. 219 (2020).

19. M. P. Colaiácovo, et al., Synaptonemal Complex Assembly in C. elegans Is Dispensable for Loading Strand-Exchange Proteins but Critical for Proper Completion of Recombination. Dev. Cell 5, 463–474 (2003).

20. S. Smolikov, et al., SYP-3 restricts synaptonemal complex assembly to bridge paired chromosome axes during meiosis in Caenorhabditis elegans. Genetics 176, 2015–2025 (2007).

21. S. Smolikov, K. Schild-Prüfert, M. P. Colaiácovo, A yeast two-hybrid screen for SYP-3 interactors identifies SYP-4, a component required for synaptonemal complex assembly and chiasma formation in Caenorhabditis elegans meiosis. PLoS Genet. 5, e1000669 (2009).

22. K. Schild-Prüfert, et al., Organization of the synaptonemal complex during meiosis in Caenorhabditis elegans. Genetics 189, 411–421 (2011).

23. S. Köhler, M. Wojcik, K. Xu, A. Dernburg, The interaction of crossover formation and the dynamic architecture of the synaptonemal complex during meiosis. bioRxiv (2020).

24. A. D. Cutter, Divergence times in Caenorhabditis and Drosophila inferred from direct estimates of the neutral mutation rate. Mol. Biol. Evol. 25, 778–786 (2008).

25. L. W. Hemmer, J. P. Blumenstiel, Holding it together: rapid evolution and positive selection in the synaptonemal complex of Drosophila. BMC Evol. Biol. 16, 91 (2016).

26. J. Fraune, et al., Hydra meiosis reveals unexpected conservation of structural synaptonemal complex proteins across metazoans. Proc. Natl. Acad. Sci. U. S. A. 109, 16588–16593 (2012).

27. J. Surkont, J. B. Pereira-Leal, Evolutionary patterns in coiled-coils. Genome Biol. Evol. 7, 545–556 (2015).

28. Z. Yang, PAML: a program package for phylogenetic analysis by maximum likelihood. Comput. Appl. Biosci. 13, 555–556 (1997).

29. S. L. Page, R. S. Hawley, The genetics and molecular biology of the synaptonemal complex. Annu. Rev. Cell Dev. Biol. 20, 525–558 (2004).

30. A. V. McDonnell, T. Jiang, A. E. Keating, B. Berger, Paircoil2: improved prediction of coiled coils from sequence. Bioinformatics 22, 356–358 (2006).

31. J. M. Dunce, et al., Structural basis of meiotic chromosome synapsis through SYCP1 self-assembly. Nat. Struct. Mol. Biol. 25, 557–569 (2018).

32. J.-F. Maure, et al., The Ndc80 loop region facilitates formation of kinetochore attachment to the dynamic microtubule plus end. Curr. Biol. 21, 207–213 (2011).

33. K.-S. Hsu, T. Toda, Ndc80 internal loop interacts with Dis1/TOG to ensure proper kinetochore-spindle attachment in fission yeast. Curr. Biol. 21, 214–220 (2011).

34. S. Yatskevich, J. Rhodes, K. Nasmyth, Organization of Chromosomal DNA by SMC Complexes. Annu. Rev. Genet. 53, 445–482 (2019).

35. M. Beasley, H. Xu, W. Warren, M. McKay, Conserved disruptions in the predicted coiled-coil domains of eukaryotic SMC complexes: implications for structure and function. Genome Res. 12, 1201–1209 (2002).

36. L. Truebestein, T. A. Leonard, Coiled-coils: The long and short of it. Bioessays 38, 903–916 (2016).

37. Z. Zhang, et al., Multivalent weak interactions between assembly units drive synaptonemal complex formation. J. Cell Biol. 219 (2020).

38. D. R. Hamill, A. F. Severson, J. C. Carter, B. Bowerman, Centrosome maturation and mitotic spindle assembly in C. elegans require SPD-5, a protein with multiple coiled-coil domains. Dev. Cell 3, 673–684 (2002).

39. S. L. Page, R. S. Hawley, c(3)G encodes a Drosophila synaptonemal complex protein. Genes Dev. 15, 3130–3143 (2001).

40. M. Sym, J. A. Engebrecht, G. S. Roeder, ZIP1 is a synaptonemal complex protein required for meiotic chromosome synapsis. Cell 72, 365–378 (1993).

41. M. J. Dobson, R. E. Pearlman, A. Karaiskakis, B. Spyropoulos, P. B. Moens, Synaptonemal complex proteins: occurrence, epitope mapping and chromosome disjunction. J. Cell Sci. 107 (Pt 10), 2749–2760 (1994).

42. R. Rillo-Bohn, et al., Analysis of meiosis in Pristionchus pacificus reveals plasticity in homolog pairing and synapsis within the nematode lineage. Cold Spring Harbor Laboratory, 662049 (2019).

43. P. M. Steinert, L. N. Marekov, R. D. Fraser, D. A. Parry, Keratin intermediate filament structure. Crosslinking studies yield quantitative information on molecular dimensions and mechanism of assembly. J. Mol. Biol. 230, 436–452 (1993).

44. A. Sato-Carlton, C. Nakamura-Tabuchi, S. K. Chartrand, T. Uchino, P. M. Carlton, Phosphorylation of the synaptonemal complex protein SYP-1 promotes meiotic chromosome segregation. J. Cell Biol. 217, 555–570 (2018).

45. R. N. McLaughlin, H. S. Malik, J. D. Levine, D. J. C. Kronauer, M. H. Dickinson, Genetic conflicts: the usual suspects and beyond. J. Exp. Biol. 220, 6–17 (2017).

46. O. Rog, S. Köhler, A. F. Dernburg, The synaptonemal complex has liquid crystalline properties and spatially regulates meiotic recombination factors. Elife 6 (2017).

47. H. B. Schmidt, D. Görlich, Nup98 FG domains from diverse species spontaneously phase-separate into particles with nuclear pore-like permselectivity. Elife 4 (2015).

48. P. Li, et al., Phase transitions in the assembly of multivalent signalling proteins. Nature 483, 336–340 (2012).

49. J. C. Newton, et al., Phase separation of the LINE-1 ORF1 protein is mediated by the N-terminus and coiled-coil domain. Biophys. J. (2021) https://doi.org/10.1016/j.bpj.2021.03.028.

50. D. M. Mitrea, R. W. Kriwacki, Phase separation in biology; functional organization of a higher order. Cell Commun. Signal. 14, 1 (2016).

51. J. B. Woodruff, et al., The Centrosome Is a Selective Condensate that Nucleates Microtubules by Concentrating Tubulin. Cell 169, 1066–1077.e10 (2017).

52. S. F. Banani, H. O. Lee, A. A. Hyman, M. K. Rosen, Biomolecular condensates: organizers of cellular biochemistry. Nat. Rev. Mol. Cell Biol. 18, 285–298 (2017).

53. A. L. Darling, Y. Liu, C. J. Oldfield, V. N. Uversky, Intrinsically disordered proteome of human membrane-less organelles. Proteomics 18, e1700193 (2018).

54. V. N. Uversky, Intrinsically disordered proteins in overcrowded milieu: Membrane-less organelles, phase separation, and intrinsic disorder. Curr. Opin. Struct. Biol. 44, 18–30 (2017).

55. C. J. Brown, et al., Evolutionary rate heterogeneity in proteins with long disordered regions. J. Mol. Evol. 55, 104–110 (2002).

56. A. Tóth-Petróczy, D. S. Tawfik, Protein insertions and deletions enabled by neutral roaming in sequence space. Mol. Biol. Evol. 30, 761–771 (2013).

57. E. V. Leushkin, G. A. Bazykin, A. S. Kondrashov, Insertions and deletions trigger adaptive walks in Drosophila proteins. Proc. Biol. Sci. 279, 3075–3082 (2012).

58. O. Podlaha, D. M. Webb, P. K. Tucker, J. Zhang, Positive Selection for Indel Substitutions in the Rodent Sperm Protein Catsper1. Mol. Biol. Evol. 22, 1845–1852 (2005).

59. O. Podlaha, J. Zhang, Positive selection on protein-length in the evolution of a primate sperm ion channel. Proc. Natl. Acad. Sci. U. S. A. 100, 12241–12246 (2003).

60. J. C. Cooper, N. Phadnis, Parallel Evolution of Sperm Hyper-Activation Ca2+ Channels. Genome Biol. Evol. 9, 1938–1949 (2017).

61. M. A. Larkin, et al., Clustal W and Clustal × version 2.0. Bioinformatics 23, 2947–2948 (2007).

62. S. Guindon, et al., New algorithms and methods to estimate maximum-likelihood phylogenies: assessing the performance of PhyML 3.0. Syst. Biol. 59, 307–321 (2010).

63. R. M. McBee, S. A. Rozmiarek, N. R. Meyerson, P. A. Rowley, S. L. Sawyer, The effect of species representation on the detection of positive selection in primate gene data sets. Mol. Biol. Evol. 32, 1091–1096 (2015).

64. M. Suyama, D. Torrents, P. Bork, PAL2NAL: robust conversion of protein sequence alignments into the corresponding codon alignments. Nucleic Acids Res. 34, W609–12 (2006).

65. L. Stevens, et al., Comparative genomics of 10 new Caenorhabditis species. Evol Lett 3, 217–236 (2019).

66. M. D. Smith, et al., Less is more: an adaptive branch-site random effects model for efficient detection of episodic diversifying selection. Mol. Biol. Evol. 32, 1342–1353 (2015).

67. D. M. Emms, S. Kelly, OrthoFinder: solving fundamental biases in whole genome comparisons dramatically improves orthogroup inference accuracy. Genome Biol. 16, 1–14 (2015).

68. D. M. Emms, S. Kelly, OrthoFinder: phylogenetic orthology inference for comparative genomics. Genome Biol. 20, 1–14 (2019).

69. S. Kumar, G. Stecher, M. Li, C. Knyaz, K. Tamura, MEGA X: Molecular Evolutionary Genetics Analysis across Computing Platforms. Mol. Biol. Evol. 35, 1547–1549 (2018).

70. S. Kumar, G. Stecher, D. Peterson, K. Tamura, MEGA-CC: computing core of molecular evolutionary genetics analysis program for automated and iterative data analysis. Bioinformatics 28, 2685–2686 (2012).

71. K. Peng, et al., Optimizing long intrinsic disorder predictors with protein evolutionary information. J. Bioinform. Comput. Biol. 3, 35–60 (2005).

72. C. Rödelsperger, et al., Spatial transcriptomics of nematodes identifies sperm cells as a source of genomic novelty and rapid evolution. Mol. Biol. Evol. (2020) https://doi.org/10.1093/molbev/msaa207.

73. C. Rödelsperger, K. Menden, V. Serobyan, H. Witte, P. Baskaran, First insights into the nature and evolution of antisense transcription in nematodes. BMC Evol. Biol. 16, 165 (2016).

74. C. Rödelsperger, et al., Phylotranscriptomics of Pristionchus Nematodes Reveals Parallel Gene Loss in Six Hermaphroditic Lineages. Curr. Biol. 28, 3123–3127.e5 (2018).

75. C. M. Phillips, K. L. McDonald, A. F. Dernburg, Cytological analysis of meiosis in Caenorhabditis elegans. Methods Mol. Biol. 558, 171–195 (2009).

76. A. Kouznetsova, R. Benavente, A. Pastink, C. Höög, Meiosis in mice without a synaptonemal complex. PLoS One 6, e28255 (2011).

77. S. E. Hitchcock-DeGregori, Tropomyosin: function follows structure. Adv. Exp. Med. Biol. 644, 60–72 (2008).

